# Mechanistic of Rac and Cdc42 synchronization at the cell edge by ARHGAP39-dependent signaling nodules and the impact on protrusion dynamics

**DOI:** 10.1101/2021.05.13.443687

**Authors:** Michael Howell, Violaine Delorme-Walker, Christopher Welch, Ritu Pathak, Klaus M. Hahn, Céline DerMardirossian

## Abstract

Compartmentalization of GTPase regulators into signaling nodules dictates the GTPase pathways selected. Rac and Cdc42 are synchronized at the cell edge for effective protrusion in motile cells but how their activity is coordinated remains elusive. Here, we discovered that ARHGAP39, a Rac and Cdc42 GTPase-activating protein, sequentially interacts with WAVE and mDia2 to control Rac/lamellipodia and Cdc42/filopodia protrusions, respectively. Mechanistically, ARHGAP39 binds to WAVE and, upon phosphorylation by Src kinase, inactivates Rac to promote Cdc42-induced filopodia formation. With our optimized FRET biosensor, we detected active Cdc42 at the filopodia tips that controls filopodia extension. ARHGAP39 is transported to filopodia tips by Myosin-X where it binds mDia2 and inactivates Cdc42 leading to filopodia retraction. Failure in lamellipodia to filopodia switch by defective ARHGAP39 impairs cell invasion and metastasis. Our study reveals that compartmentalization of ARHGAP39 within Rac/Cdc42 signaling nodules orchestrates the synchronization of lamellipodia/filopodia crosstalk and highlights the intricate regulation of leading edge dynamics in migrating cells.

## INTRODUCTION

Protein-protein interactions are critical for cellular signal transductions that play key roles in both normal and aberrant functions in cells. Changes in specificity and affinity of these interactions can lead to pathological conditions, such as uncontrolled cell growth and/or cell migration. Many cellular responses require the coordination of the precise integration of localized, transient signalling events with the remodelling of the actin cytoskeleton. The small GTPases Rac and Cdc42 are spatiotemporally synchronized at the cell leading edge (Machacek et al., 2009) to drive cytoskeleton dynamics and thus regulate the cell edge protrusions and migration. GEFs (Guanine Exchange Factors) and GAPs (GTPases activating proteins) outnumber the GTPases, reflecting the complex and context-specific mechanisms of activation. Compartmentalization of Rac and Cdc42, their regulators and their effectors in one key strategy used by the cellular machinery to achieve signal specificity (Etienne-Manneville and Hall, 2002). GAP proteins contain multifunctional domain features (Moon and Zheng, 2003; Tcherkezian and Lamarche-Vane, 2007) that might select specific signaling pathways in time and space. However, the molecules that tether GAPs to restrict their activity spatially and temporally towards specific GTPases are challenging to be found. ARHGAP39 is ubiquitously expressed (Nowak, 2018) and, in addition to a GAP catalytic domain, contains two WW domains (Lundstrom et al., 2004), known to mediate protein-protein interactions (Kay et al., 2000). The ARHGAP39 homologue in Drosophila, Vilse, has been characterized as a GAP for Rac and Cdc42 (Lim et al., 2014; Lundstrom et al., 2004) and its WW domains binds to the Slit receptor, Robo, to mediate the axon guidance repulsion during cell migration (Hu et al., 2005; Lundstrom et al., 2004). Most of the findings with ARHGAP39/Vilse have been made in the brain. More recently, bioinformatic analyses have revealed the increase of expression level or mutations associated with many types of cancer (cancer.sanger.ac.uk/cosmic/gene). Here, we identified WAVE and mDia2 that bind and compartmentalize the activity of ARHGAP39 toward Rac and Cdc42, respectively, to coordinate lamellipodia and filopodia protrusions, and to regulate filopodia dynamics in migrating cancer cells.

The WAVE family of proteins drive membrane protrusion, lamellipodia, by coordinating upstream signals and generating short, highly branched actin networks *via* activation of the Arp2/3 complex (Chen et al., 2014; Lebensohn and Kirschner, 2009). WAVE is part of the WAVE regulatory complex (WRC) and is recruited to the plasma membrane by active Rac (Chen et al., 2010; Steffen et al., 2004). Deciphering the molecular machineries that regulate Rac activation and, thus, WAVE activity is crucial for understanding how the active WRC is controlled at the membrane and how filopodia are generated. Filopodia are thin cellular filaments that dynamically extend and retract to act as environmental sensors for migrating cells (Davenport et al., 1993). The Diaphanous-related formin mDia2 is required for filopodia extension. mDia2 generates long unbranched actin filaments (Pruyne et al., 2002) on which a motor molecule Myosin-X (Myo10) transports proteins to regulate filopodia function (Tokuo and Ikebe, 2004; Zhang et al., 2004). At the plasma membrane, mDia2 is associated with the WRC. Activation of mDia2 is dependent on both WAVE removal from the membrane (Beli et al., 2008) and the binding of active Cdc42 (Alberts, 2001; Alberts et al., 1998). Once active, mDia2 induces the assembly of actin filaments by processive elongation in the growing filopodia (Kovar et al., 2006; Schirenbeck et al., 2005) and remains at the filopodia tips.

Although WAVE and mDia2 are regulated by active Rac and Cdc42, respectively, the mechanisms that control the activation state of both GTPases have remained unknown. Furthermore, the mechanism that controls the inactivation of mDia2 in the filopodia and therefore modulates filopodia extension is not clear. Here, we address how lamellipodia signalling is coordinated with pathways regulating filopodia dynamics to control cell edge protrusion and thus, cell migration and invasion. We identified a highly sophisticated Rac and Cdc42 signaling network regulated by ARHGAP39. Together, our results provide a comprehensive mechanistic understanding of how ARHGAP39 controls the cell leading edge and cell migration.

## RESULTS

### ARHGAP39 directly interacts and co-localizes with WAVE in a WW domain-dependent fashion

In an effort to better understand cell migration, we identified ARHGAP39 after surveying the genome for uncharacterized GAPs containing WW domains, known to interact with polyproline regions, as proline-rich motifs are common to cytoskeletal proteins (Kay et al., 2000). Using a pull-down GTPase activity assay in ARHGAP39-depleted cells (Figures S1A and S1B), we confirmed that ARHGAP39 is a GAP for Rac and Cdc42 but not for RhoA (Figures 1A and 1B). To next investigate the function of ARHGAP39, we sought to identify proteins that interact with its WW domains by mass spectrometry and pull-down experiments. We found an interaction with both Nck-associated protein 1 (Nap1), a member of the WRC (Chen et al., 2017), as well as Drf3/mDia2 (hereafter mDia2). The presence of Nap1 suggested that ARHGAP39 might associate with the WRC and interact with WAVE, which contains polyproline sequences that match the consensus binding motifs for the WW domains in ARHGAP39 (Ingham et al., 2005). The existence of a complex was assessed by pull-down assays which showed that ARHGAP39-Wdom, consisting of the WW domains but not the GAP domain, directly interacted with WAVE and specifically bound the WAVE polyproline region (PP) (Figures 1C and 1D). We further confirmed that endogenous WAVE from cell lysates interacted with Wdom (Figures S1C and S1D). We next observed the interaction between ARHGAP39 and WAVE in cells by immunofluorescence analysis. ARHGAP39 wild-type (WT) did not localize at the plasma membrane under basal conditions (Figure 1E, top panels, and 1F, white columns). However, co-expression with constitutively active Rac (Rac Q61L) resulted in ARHGAP39-WT translocation to the membrane (Figure 1E, bottom panels, and 1F, black columns) where it co-localized with endogenous WAVE (Figures 1E, 1G and Figures S1E and S1F). We further validated the specificity of the WAVE interaction by overexpressing an ARHGAP39 WW domain mutant (ARHGAP39-Wmut) incapable of binding to polyproline ligands. ARHGAP39-Wmut failed to co-localize with endogenous WAVE even in the presence of constitutively active Rac (Figures 1 E-G and S1E-F). We next tested if this event was dependent on ARHGAP39 GAP activity. ARHGAP39-R963A (RA), a point mutation of the conserved Arg residue within the GAP domain, known to render a GAP protein inactive towards GTPase substrates (Barrett et al., 1997), increased at the membrane in the presence of active Rac and co-localized with WAVE, as did ARHGAP39-Wdom, indicating that ARHGAP39 recruitment did not depend on a functional GAP domain (Figures 1E-G and S1E-F). Altogether, these results show that ARHGAP39 binds to and co-localizes with WAVE at the membrane when it is part of the active WRC, and this interaction relies on ARHGAP39 WW domains.

**Figure 1.**
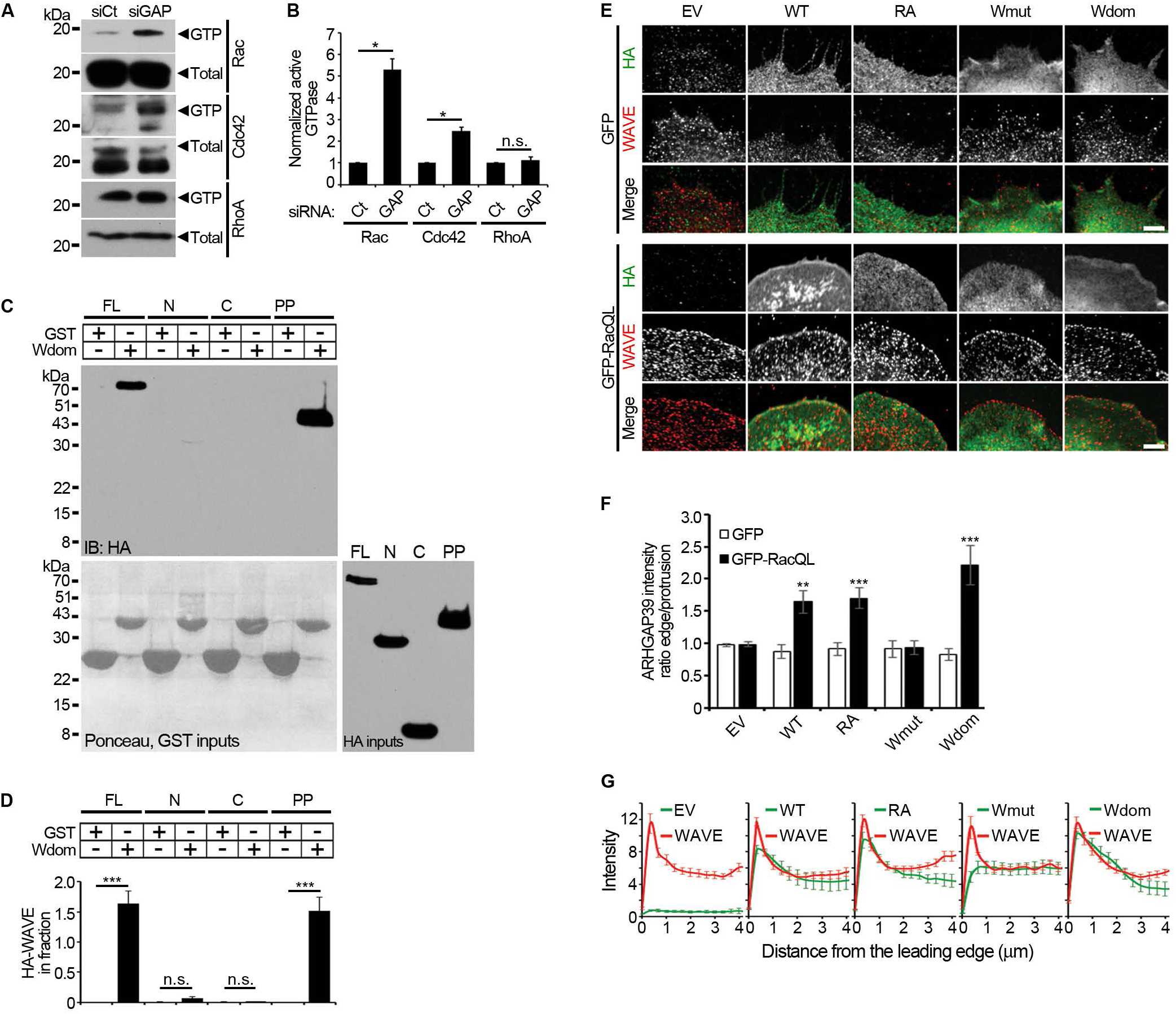
ARHGAP39 is a GAP for Rac and Cdc42 and is recruited to the cell edge by active Rac. (A and B) ARHGAP39 GTPase assay. (A) Lysates of HeLa cells transfected with control or ARHGAP39 siRNA oligos for 48 h were subjected to a GST-PBD or -RBD GTPase binding assay to pull down active Rac/Cdc42 or RhoA, respectively. (B) Quantification of ARHGAP39 GTPase activity. (C and D) ARHGAP39 N-terminal domain directly interacts with WAVE polyproline region. (C) In vitro translated HA-WAVE constructs full-length (FL), N-terminal (N), C-terminal (C) and poly-proline rich domain (PP) were incubated with GST or GST-ARHGAP39 aa1-134 (Wdom). WAVE proteins were detected using anti-HA antibodies. Bottom left panel, red ponceau stain of GST inputs. Bottom right panel, HA-WAVE inputs. (D) Quantification of WAVE binding to ARHGAP39. (E-G) Active Rac recruits ARHGAP39 to the cell leading edge. (E) HeLa cells were transfected with GFP or GFP-RacQ61L in combination with HA empty vector (EV), HA-ARHGAP39-WT, HA-ARHGAP39-RA (GAP inactive), HA-ARHGAP39-Wmut (WAVE-binding deficient) or HA-Wdom (ARHGAP39 aa 1-134). Cells were labelled with anti-HA and anti-WAVE antibodies to visualize ARHGAP39 (green) and endogenous WAVE (red), respectively. Scale bars, 5 µm. (F) Average fluorescence intensity ratio of ARHGAP39 at the cell edge *vs* 3 µm inside the protrusion. *n* ≥ 60 cells for each condition. (G) ARHGAP39 co-localizes with WAVE at the cell edge. Average fluorescence intensity of HA (green) and WAVE (red) from cells expressing RacQL. Intensity profiles start at the cell leading edge (0 µm) and end at 4 µm inside the protrusion. n ≥ 20 cells for each condition. Data are representative of three independent experiments and are presented as mean ± SEM. Statistical analysis was performed using two-tailed Student’s *t*-test. n.s., not statistically significant; *\**p < 0.05; *\*\**p < 0.01 and ***p < 0.001.

### Interaction with WAVE and phosphorylation of residues Y313,325 are critical for the GAP activity of ARHGAP39

Phosphorylation is one mechanism commonly used by cells to regulate GAPs (Tcherkezian and Lamarche-Vane, 2007). ARHGAP39 tyrosine residues 313, 325 and 785, are contained in potential Src phosphorylation consensus motifs (EEIYGEFD/F) (Songyang et al., 1995) and are conserved across human, mouse, chicken or zebrafish (Figure 2A). Kinase assays confirmed that ARHGAP39-WT was phosphorylated by Src (Figures 2A and 2B). The single mutation of Y313, Y325, or Y785 to alanine or double mutations Y313,325A (2YA) each decreased ARHGAP39 phosphorylation, whereas the combined alanine mutations (3YA) completely abrogated ARHGAP39 phosphorylation. We next examined whether WAVE binding was required for ARHGAP39 phosphorylation and found that ARHGAP39-Wmut was phosphorylated by Src. However, we determined that ARHGAP39-Wmut Y785A phosphorylation was markedly decreased in contrast to ARHGAP39-Wmut, Y313/325A, Y313A or Y325A (Figures 2C and 2D), indicating that only Y785 was phosphorylated in ARHGAP39-Wmut. We analyzed whether ARHGAP39 phosphorylation was required for its recruitment at the membrane.

**Figure 2.**
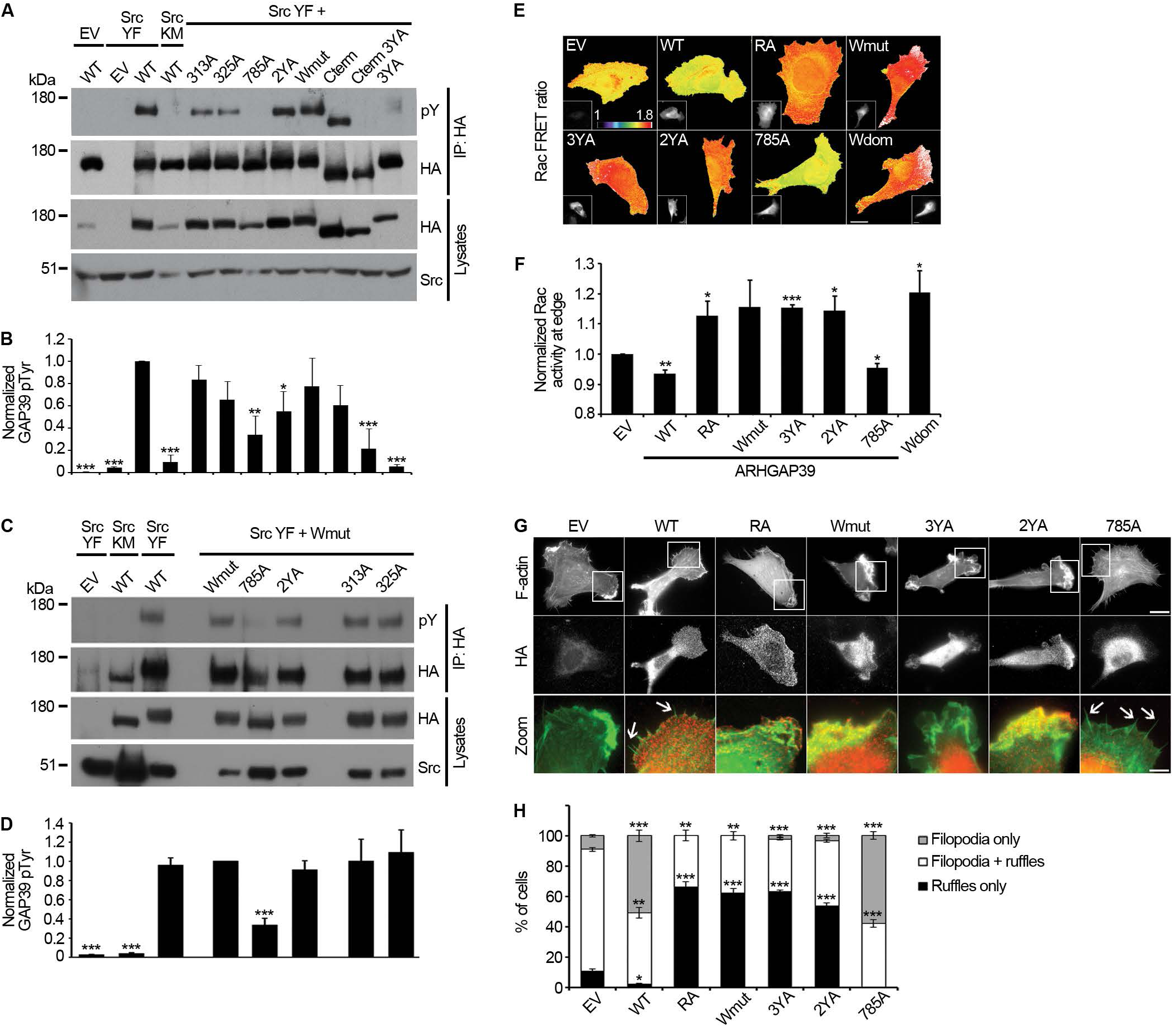
ARHGAP39 activity is regulated by both phosphorylation and binding to WAVE. (A and B) ARHGAP39 is tyrosine phosphorylated by Src kinase. (A) ARHGAP39 was immunoprecipitated from HeLa cells co-expressing constitutively active or inactive Src kinase (Src YF or Src KM, respectively) and HA-ARHGAP39-WT, −313A, −325A, −785A, -Wmutant, -Cterm, -Cterm 3YA or -3YA mutants. Tyrosine phosphorylation, ARHGAP39 and Src were detected using anti-phosphotyrosine (pY), -HA and -Src antibodies, respectively. (B) Quantification of ARHGAP39 tyrosine phosphorylation. (C and D) ARHGAP39 Wmutant is phosphorylated on residue Y785. (C) ARHGAP39 was immunoprecipitated from HeLa cells co-expressing Src YF or Src KM and HA-ARHGAP39 -WT, - Wmutant, -Wmutant 785A, -Wmutant 2YA, -Wmutant 313A or –Wmutant 325A. Tyrosine phosphorylation, ARHGAP39 and Src were detected using anti-phosphotyrosine (pY), -HA and -Src antibodies, respectively. (D) Quantification of ARHGAP39 tyrosine phosphorylation. (E and F) Ratiometric images of HeLa cells (E) transfected with Rac1 biosensor in combination with HA empty vector (EV), HA-ARHGAP39 -WT, -RA, -Wmut, −3YA, −2YA, −785A or -Wdom. Insets show HA expression. Scale bars, 15 μm. (F) Normalized Rac1 activity 0 to 3 μm from the leading edge. n = 27-53 cells. (G and H) HT-1080 cells expressing either HA empty vector (EV), HA-ARHGAP39 -WT, -RA, -Wmut, −3YA, −2YA, or −785A were labelled with phalloidin and anti-HA antibodies to detect F-actin (green) and ARHGAP39 (red), respectively (G). Scale bar, 15 μm. White boxes in F-actin indicate regions used for magnified images in lower panel. Scale bar, 5 μm. White arrows indicate filopodia. (H) Average percentage of cells with ruffles only (black), mixed filopodia and ruffles (white) or filopodia only (grey). n = 63-98 cells for each condition. Data are representative of three to six (A,B), four (C,D) or three (E-H) independent experiments and are presented as mean ± SEM. Statistical analysis was performed using two-tailed Student’s *t*-test. *\**p < 0.05; *\*\**p < 0.01 and *\*\*\**p < 0.001.

ARHGAP39-WT or the different non-phosphorylatable mutants all translocated to the membrane and co-localized with WAVE in cells expressing constitutively active Rac (Figures 2B and 2C). Taken together, these results indicate that Src phosphorylation of residues Y313 and Y325, but not Y785, occurs after ARHGAP39 binding to WAVE at the membrane.

We next investigated whether ARHGAP39 phosphorylation modulated its GAP activity in cells using a Rac biosensor (Miskolci et al., 2016). ARHGAP39-WT expression significantly decreased Rac activation, whereas inactive GAP constructs (RA or Wdom) abrogated ARHGAP39 activity on Rac (Figures 2E and 2F). With an efficiency similar to inactive GAP, ARHGAP39-3YA or 2YA could not decrease the level of active Rac while ARHGAP39-Y785A still had functional GAP activity. Finally, we determined that the association with WAVE was necessary for ARHGAP39 GAP function, as cells expressing ARHGAP39-Wmut showed Rac activity similar to cells expressing inactive ARHGAP39 proteins (Figures 2E and 2F). These results demonstrate that both the binding to WAVE and the phosphorylation by Src on two residues Y313 and Y325, but not Y785, are critical for promoting ARHGAP39 catalytic activity.

Although Rac-dependent activation of WAVE induces the formation of lamellipodia, the inactivation of Rac, and thus WAVE removal from the membrane, is a necessary step for filopodia generation (Beli et al., 2008). We next investigated whether ARHGAP39 activity on Rac supported filopodia formation. In control cells, the majority of cells exhibited a mix of both membrane ruffles, indicative of lamellipodia and active Rac, and filopodia, associated with Rac inactivation (Figures 2G and 2H, white column) and approximately 10% of cells showed the presence of either filopodia (Figure 2H, gray column) or membrane ruffles (Figure 2H, black column). Expression of ARHGAP39-WT markedly increased the number of cells with filopodia (Figures 2G and 2H, gray column). Conversely, ARHGAP39 constructs that could not inactivate Rac (RA, Wmut or 3YA) strongly increased the number of cells with membrane ruffles (Figures 2G and 2H, black column). ARHGAP39-Y785A promoted the formation of filopodia with efficiency similar to ARHGAP39-WT (Figures 2G and 2H, gray column), confirming that this residue is not involved in ARHGAP39 activity. Altogether, these data indicated that both the integration of ARHGAP39 into the active WRC via the binding to WAVE, and its GAP activity towards Rac, are essential steps for ARHGAP39 contribution to filopodia formation.

### Tyrosine 785 phosphorylation is required for ARHGAP39 binding to Myo10 and its localization within filopodia

While investigating the effect of ARHGAP39 on filopodia formation, we observed that ARHGAP39-WT localized in the filopodia shaft and was enriched at some filopodia tips (Figures 3A, white arrow, and 3B), suggesting that ARHGAP39 might also function as a regulator of filopodia dynamics. To test this hypothesis, we first investigated the mechanism of ARHGAP39 localization within filopodia. To generate filopodia formation by bypassing the Rac/WAVE pathway, we expressed constitutively active mDia2 in cells, using truncated mDia2 (mDia2ΔDAD) as full-length mDia2 is conformationally closed (Alberts, 2001; Otomo et al., 2005; Peng et al., 2003). ARHGAP39-Y313A, Y325A or 2YA still localized within filopodia (Figure 3C). However, ARHGAP39-Y785A or ARHGAP39-3YA failed to enter filopodia, indicating that Y785 phosphorylation is critical for the presence of ARHGAP39 within filopodia. The detection of mDia2 in the mass spectrometry pull-down assay with ARHGAP39-Wdom prompted us to examine whether mDia2 played a direct role in ARHGAP39 filopodia localization. Pull-down experiments determined that ARHGAP39-Wdom directly bound to active mDia2ΔDAD and showed a strong interaction with the mDia2 proline-rich FH1 region (Figures S3A and S3B). Immunofluorescence analysis indicated that ARHGAP39-WT and mDia2ΔDAD co-localized at the filopodia tip, but not in every filopodia and not in the filopodium shaft (Figures S3C and S3D). We further examined whether mDia2 was required for ARHGAP39 localization within filopodia by depleting cells of mDia2. Because the absence of mDia2 abrogated active Cdc42-induced filopodia formation (Figures S3E and S3F), cells were transfected with Myo10, known to initiate filopodia formation (Berg and Cheney, 2002). ARHGAP39 not only still localized to filopodia in mDia2-depleted cells, but its presence was also increased at filopodia tips (Figures S3G and S3H). Notably, Myo10 and ARHGAP39 co-localized in the shaft and at the tip of filopodia (Figures 3D and 3E). Using pull-down assays, we found that ARHGAP39-WT or ARHGAP39-2YA interacted with Myo10, but not ARHGAP39-Y785A or ARHGAP39-3YA (Figures 3F and 3G). We next monitored the dynamics of ARHGAP39 within filopodia by time-lapse imaging. We observed that accumulation of ARHGAP39 at the filopodia tip preceded filopodia retraction (Figure 3H, white arrow at 30s; Supplementary Video 1). These data demonstrate that phosphorylation on residue Y785 by Src kinase is critical for the transport of ARHGAP39 by Myo10 within filopodia and suggest that ARHGAP39 at the tip of filopodia controls filopodia extension/retraction.

**Figure 3.**
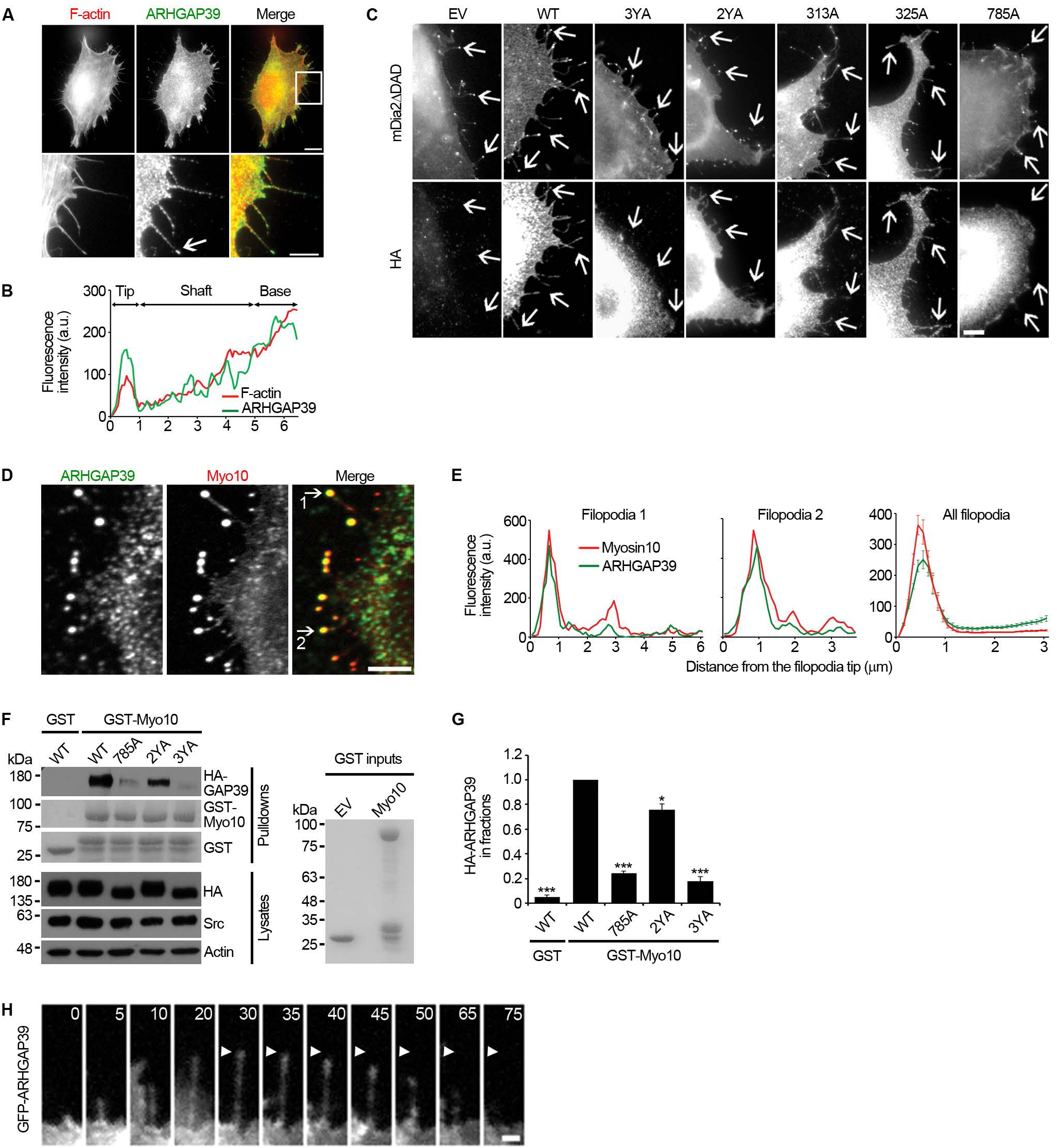
Y785-phosphorylated ARHGAP39 is transported by Myo10 in filopodia. (A and B) ARHGAP39 localizes to filopodia. (A) Image of a HeLa cell expressing HA-ARHGAP39 WT (green) and labelled for F-actin (red). Scale bar, 10 μm. White box indicates the region used for magnification in bottom panel. Scale bar, 5 μm. White arrow indicates the filopodia used for the line scan in (B). (B) Intensity profiles of ARHGAP39 (green) and F-actin (red) starting at the tip of the filopodia (0 μm) and ending at the protrusion (6 μm). (C) Y785 phosphorylation is required for ARHGAP39 localization to filopodia. HeLa cells were transfected with GFP-mDia2ΔDAD in combination with either HA empty vector (EV), HA-ARHGAP39 - WT, −3YA, −2YA, −313A, −325A or −785A. Cells were labelled with anti-HA antibodies to detect ARHGAP39. Arrows indicate filopodia. Note the absence of ARHGAP39 −3YA or −785A in filopodia. (D and E) ARHGAP39 co-localizes with Myo10 in the filopodia. (D) Cells depleted in mDia2 were co-transfected with GFP-Myo10 (red) and HA-ARHGAP39 WT (green). Cells were labelled with anti-HA antibodies to detect ARHGAP39. Scale bar, 5 μm. White arrows indicate two representative filopodia used for line scans in (E). (E) Intensity profiles of ARHAGP39 (green) and Myo10 (red) starting at the tip of the filopodia. Most right panel shows the average intensity profiles of 100 filopodia. (F and G) Y785 phosphorylation is required for ARHGAP39 binding to Myo10. (F) Lysates of HeLa cells co-expressing SrcYF and HA-ARHGAP39 -WT, −785A, −2YA or −3YA were incubated with GST- or GST-Myo10 Myth-FERM. ARHGAP39 was detected using anti-HA antibodies. Right panel, red ponceau stain of GST inputs. (G) Quantification of ARHGAP39 binding to Myo10 Myth-FERM domain normalized to ARHGAP39 levels in cell lysates and expressed relative to ARHGAP39-WT binding to Myo10. Statistical analysis was performed using two-tailed Student’s t-test. *\**p < 0.05 and *\*\*\**p < 0.001 compared to WT. (H) Time course imaging of GFP-ARHGAP39 dynamics in filopodia. Time is shown in seconds. Accumulation of ARHGAP39 at tip (white arrowhead) precedes filopodia retraction. Scale bar, 1 μm. Data are representative of three independent experiments and are presented as mean ± SEM.

### ARHGAP39 controls Cdc42 inactivation and filopodia dynamics

Our findings encouraged us to determine the molecular mechanism through which ARHGAP39 regulates filopodia retraction. The discovery of a GAP for Cdc42 that localizes to filopodia and binds mDia2 (Figure S3A) suggested that active Cdc42 is present within filopodia and that ARHGAP39 controls filopodia dynamics by regulating Cdc42 activity within filopodia. Although several studies have implicated Cdc42 in the initial formation of filopodia (Lammers et al., 2008; Pruyne et al., 2002; Yasuda et al., 2004), the localization and activity of Cdc42 in filopodia have remained unclear. To investigate Cdc42 activation status within filopodia, we developed an optimized Cdc42 biosensor based on our earlier designs (Machacek et al., 2009; Nalbant et al., 2004; Pertz et al., 2006) and verified its overall functionality (Figures S4A and S4B). Cdc42 activity was then detected in live cells (Figure 4A and Supplementary Video 2). Quantification of the FRET ratio showed active Cdc42 at the filopodia tip (Figure 4B). If active Cdc42 must remain bound to mDia2 to keep it in an active conformation, it follows that Cdc42 inhibition and detachment from mDia2 would inactivate the complex. Because mDia2 controls filopodia growth, the activation of Cdc42 should be lower when filopodia are retracting. To test this hypothesis, we analyzed Cdc42 activity during the entire filopodial extension/retraction cycles (Figure 4C and Supplementary Video 3), discerned by elongation then shortening of the same filopodium. First, we found that in membrane regions where the formation of a single filopodium was discernible, a highly spatially restricted burst of Cdc42 activation marked the area where filopodia formed prior to filopodia elongation (Figures 4C and 4D). Second, as the filopodia grew, the pool of active Cdc42 was carried upward from the membrane into the filopodia tip (Figure 4C, white arrow). Cdc42 showed an average higher FRET signal when filopodia were in the extension phase than in the retraction phase (Figures 4C and 4E). Moreover, we observed that a low level of active Cdc42 at the tip preceded filopodia retraction (Figure 4C, red arrow). These data suggest that active Cdc42 controls both filopodia formation and extension, whereas its inactivation leads to filopodia retraction.

**Figure 4.**
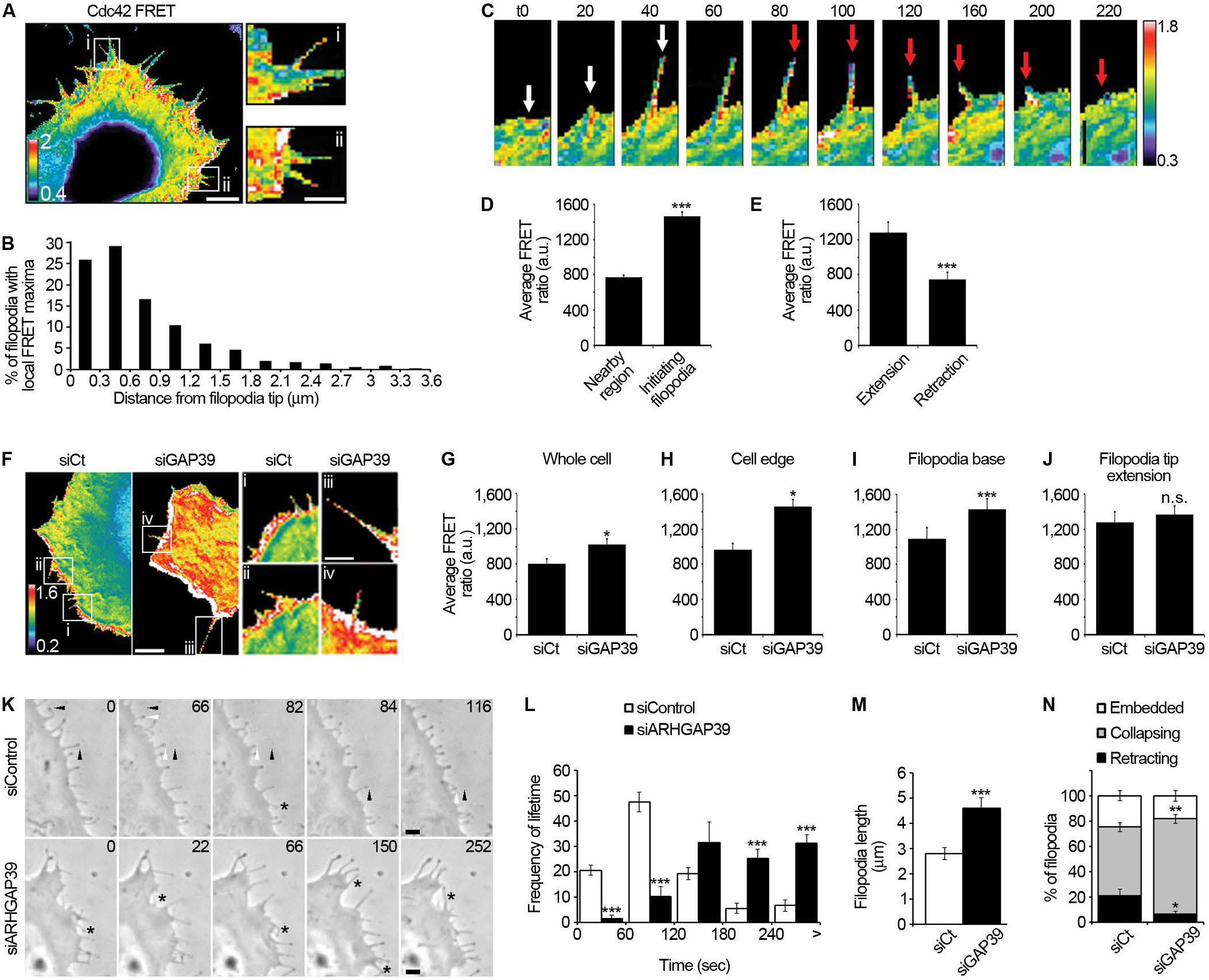
Endogenous ARHGAP39 regulates Cdc42 activity in filopodia and controls filopodia dynamics. (A-E) Cdc42 activation in filopodia correlates with filament dynamics. (A) Cdc42 activation is detected at the leading edge and in filopodia. Processed FRET images of a Hela cell transfected with the Cdc42 biosensor. Scale bar, 5 μm. Scale bar in close up panels, 2.5 μm. (B) Cdc42 is active at filopodia tips. Positioning of the most common maxima of Cdc42 signal along the filopodia length as measured from the tip. n = 345 filopodia. Cdc42 FRET average maximum distance is 0.972 μm from distal end. (C) High FRET ratio (Cdc42 activation) precedes filament formation and elongation (white arrow) while low Cdc42 activity precedes filament retraction (red arrow). Time is shown in seconds. Scale bar, 3 μm. (D) Cdc42 activation occurs at the cell periphery prior to filopodium formation. Quantification of FRET ratio at initiating filopodia compared to a nearby region. n = 46 filopodia. *\*\*\**p < 10^−21^. (E) Cdc42 activation is higher during filopodia extension versus retraction. n = 50 filopodia extending and 30 filopodia retracting. *\*\*\**p < 10^−17^. (F-J) Cdc42 activity is regulated by ARHGAP39. (F) Processed FRET images of HeLa cells treated with siRNA control (left panel) or siRNA ARHGAP39 (right panel). Scale bar, 8 μm. Scale bar in close up panels, 4 μm. Quantification of Cdc42 activation in siRNA control or siRNA ARHGAP39 treated cells: (G) whole cell and (H) cell edge (region 0-2 μm from cell leading edge). n = 6 and n = 7 cells for siControl and siARHGAP39, respectively. (I) filopodia base, measured during filopodia growth. n = 62 and n = 38 filopodia for siCt and siGAP39, respectively. *\*\*\**p < 10^−8^; (J) filopodia tips during extension. n = 50 and n = 34 filopodia for siControl and siARHGAP39, respectively. (K-N) ARHGAP39 depletion affects filopodia dynamics. (K) Brightfield time course imaging of filopodia dynamics in HeLa cells transfected with control or ARHGAP39 siRNA oligos in combination with HA-Myo10. Time is shown in seconds. Black arrowheads indicate filopodia tip before retraction and white arrowheads show the same filopodia tip after retraction. Black stars indicate collapsing filopodia. Scale bars, 2 μm. (L) Quantification of the percentage of filopodia retracting (black), collapsing on membrane (grey) or embedded in membrane (white) in siControl and siARHGAP39 cells. *n* = 436 and 225 filopodia, respectively. (M) Quantification of the frequency of filopodia lifetime in siControl (white) and siARHGAP39 (black) cells. *n* = 513 and 147 filopodia, respectively. (N) Average filopodia length in siControl (white) and siARHGAP39 (black) cells. *n* = 691 and 364 filopodia, respectively. Data are representative of three (A-J) or two (K-N) independent experiments and are presented as mean ± SEM. Statistical analysis was performed using two-tailed Student’s *t*-test. n.s., not statistically significant; *\**p < 0.05; *\*\**p < 0.01 and *\*\*\**p < 0.001 unless specified.

We next characterized the function of ARHGAP39 on Cdc42 within filopodia. We found that ARHGAP39 depletion markedly decreased the number of filopodia formed per cell compared to siRNA control (Figures S4C and S4D). Expression of siRNA-resistant ARHGAP39 protein not only reversed this phenotype but also significantly increased the number of filopodia per cell relative to control cells. This result reinforced our findings that suppression of Rac activity by ARHGAP39 is required for the formation of filopodia (Figures 2E-H). Compared to control cells, Cdc42 activity was significantly increased in ARHGAP39-depleted cells, throughout the cell (Figures 4F and 4G and Supplementary Video 4), at the cell edge (Figure 4H), at the filopodia base prior to filopodium formation (Figure S4E), and during filopodia growth (Figure 4I). These foci likely contain newly activated mDia2, as the pool of active Cdc42 is carried upward from the membrane into the filopodia tip. The level of active Cdc42 level was not affected at the tip of elongating filopodia relative to the control (Figure 4J). Live-cell imaging of ARHGAP39-depleted cells showed a significant increase in filopodia lifetime (Figures 4K and 4I, black column and Supplementary Videos 5 and 6) and in filopodia length (Figure 4M, black column) compared to control cells (Figure 4L and 4M, white column). We found that the number of filopodia embedded within the membrane was not impaired in ARHGAP39-depleted cells (Figure 4N, white column) but the number of retracting filopodia was significantly decreased (Figure 4N, black column) while the percentage of filopodia that collapsed was markedly increased (Figure 4N, grey column) compared to control siRNA cells. Altogether, these results suggest that mDia2 activity is regulated by ARHGAP39 in filopodia and that ARHGAP39 is a critical factor in maintaining the filopodia extension/retraction equilibrium.

### ARHGAP39 regulates cancer cell motile and invasive properties

Having dissected the molecular mechanism of ARHGAP39 function at the cell leading edge, we next investigated the biological impact of ARHGAP39 on cell migration and invasion. ARHGAP39 depletion remarkably decreased cell migration in 3D collagen gels compared to control cells (Figure 5A and Supplementary Videos 7 and 8); the average of both total and net distances (Figure 5B), directionality (Figure 5C) as well as velocity (Figure 5D) were significantly below control levels. We further examined whether ARHGAP39 might play a role in the invasion steps during the metastatic process by evaluating its effect on intravasation and extravasation in vitro. ARHGAP39 ablation or ARHGAP39 mutants deficient in the Rac/WAVE pathway, either GAP inactive (RA, 3YA, Wdom) or WAVE-binding deficient (Wmut), markedly impaired both the intercalation (Figures S5A and S5B) and the migration of cancer cells through endothelial cell monolayers compared to control (Figures 5E and 5F). As these conditions all prevented the formation of filopodia, we next determined whether the function of ARHGAP39 in filopodia alone is sufficient to control cancer cell invasion. We examined the invasive properties of cancer cells expressing ARHGAP39-Y785A, a mutant that inactivates Rac, generates the formation of filopodia, but cannot regulate filopodia dynamics due to its absence within filopodia. Intravasation (Figures 5E and 5F) and extravasation (Figures S5A and S5B) of cancer cells were both significantly reduced by the expression of ARHGAP39-Y785A. Among the few cells that intercalated in the endothelial monolayer, we found that these cells presented an exaggerated number of protrusions and an increased cell thickness indicative of a failure to properly integrate the monolayer (Figures S5C-E). Taken together, these results indicate that ARHGAP39 regulation of both lamellipodia/filopodia coordination and filopodia dynamics is essential to cancer cell migration and invasion.

**Figure 5.**
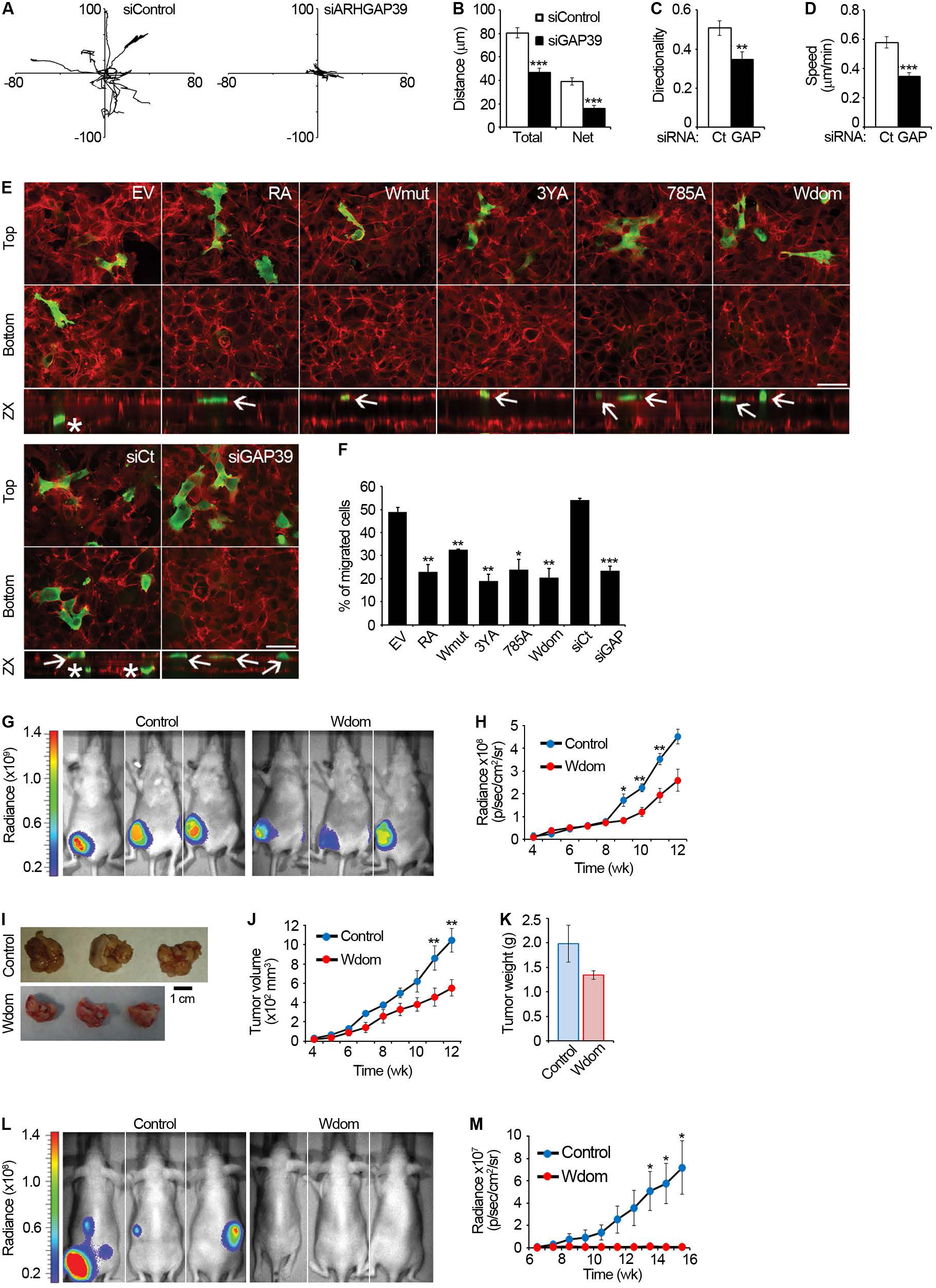
ARHGAP39 regulates cancer cell motile and invasive properties. (A-D) ARHGAP39 depletion decreases cancer cell migration in 3D. (A) Individual tracks of ten cells transfected with control or ARHGAP39 siRNA oligos transposed to a common origin. Quantification of motility parameters, including total and net distances (B), directionality (C) and cell velocity (D). n = 35 cells for each condition. (E and F) ARHGAP39 regulates cancer cell trans-endothelial migration. (E) Images of the top and bottom of transwells after trans-endothelial migration assay with HT-1080 cells transfected with GFP empty vector (EV), HA-ARHGAP39 -WT, -RA, -Wmut, −3YA, −785A, -Wdom, siRNA control or siRNA ARHGAP39. Cells were labelled with anti-HA antibodies and phalloidin to detect ARHGAP39 (green) and F-actin (red), respectively. Scale bar, 50 μm. Bottom panels are confocal z projections of entire transwell. White asterisks indicate cells that have migrated through the endothelial cell monolayer. White arrows indicate cells that did not migrate. (F) Quantification of the percentage of cells intravasating for each condition. n = 46 to 94 cells. (G-K) Wdom expression reduces mammary tumor growth. (G) Noninvasive bioluminescence imaging 10 weeks after mammary fat pad injection of 5×10^5^ control or Wdom-expressing stable MDA-MB-435 cells (n = 5 mice). (H) Quantification of average radiance over time. (I) Mammary fat pads of mice injected with control or Wdom stable MDA-MB-435 cells, 12 weeks post-injection. (J) Average tumor size measured over time with a caliper. (K) Average tumor weight of removed mammary fat pads. (L and M) Wdom expression inhibits tumor formation from the bloodstream. (L) Noninvasive bioluminescence imaging 15 weeks after i.v injection of 5×10^5^ control or Wdom-expressing stable MDA-MB-435 cells (n = 5 mice). (M) Quantification of average radiance over time. Data are representative of three (A-F) independent experiments. Data are presented as mean ± SEM. Statistical analysis was performed using two-tailed Student’s *t*-test. *\**p < 0.05; *\*\**p < 0.01 and \**\*\**p < 0.001.

We next determined whether preventing the lamellipodia/filopodia synchronization affected tumor development and invasion in mice. We developed stable cancer cells carrying control or ARHGAP39-Wdom and verified that the latter were inhibited in their invasive capacities as observed in vitro (Figures S5F and S5G). In a xenograft tumor model, both control and ARHGAP39-Wdom stable cell lines developed tumors in mice mammary pads (Figure 5G). However, the tumor growth rate of ARHGAP39-Wdom xenografts was slower (Figures 5H and 5J) and the tumors were smaller (Figures 5I and 5K). Cell division in 2D (Figure S6A) or in 3D soft agar (Figures S6B-E) was not altered by the expression of ARHGAP39-Wdom, suggesting that the tumor development observed in mice might be affected by the surrounding microenvironment. In an experimental metastasis model, mice injected with control cancer cells developed a tumor that grew regularly over 16 weeks, whereas ARHGAP39-Wdom expression inhibited tumor development (Figures 5L and 5M), indicating a defect in cell invasion from the bloodstream through the endothelial cells. Collectively, these results support an oncogenic role for ARHGAP39 and demonstrate that ARHGAP39-dependent coordination of lamellipodia to filopodia is critical for cancer cell invasion.

In sum, our findings suggest the following model: ARHGAP39 binds WAVE at the plasma membrane. Following ARHGAP39 phosphorylation on Y313 and Y325 by Src, Rac is inactivated and mDia2 initiates the formation of filopodia. Myo10 carries ARHGAP39 phosphorylated on Y785 to filopodia tips. There, ARHGAP39 binds active mDia2, regulates the activation of Cdc42 interacting with mDia2, and ultimately controls filopodia dynamics (Figure 6).

**Figure 6.**
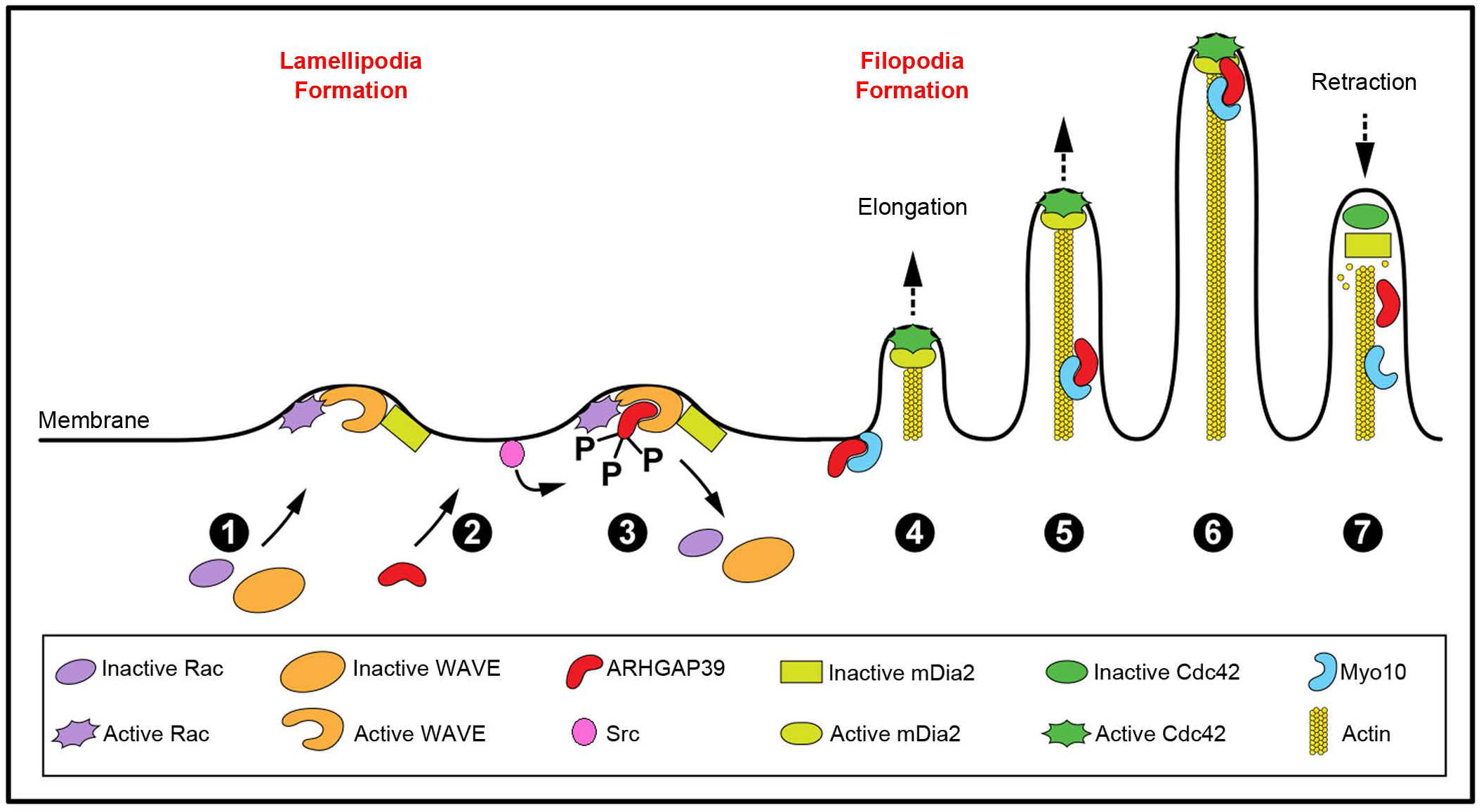
Graphical model of how ARHGAP39 regulates Rac/lamellipodia and Cdc42/filopodia protrusions in cells. (1) Inactive cytosolic Rac translocates to the plasma membrane where it is activated. In turn, Rac activates the WAVE complex that drives the formation of lamellipodia(Chen et al., 2010; Steffen et al., 2004). (2) Cytosolic ARHGAP39 is then recruited at the cell edge by active Rac. (3) Both the ARHGAP39 binding to WAVE and its phosphorylation by Src are required for its GAP activity towards Rac. Once Rac is inactivated, WAVE dissociates from the plasma membrane. (4) mDia2 freed from the WAVE complex(Beli et al., 2008) binds to active Cdc42 and induces the formation of filopodia. (5) ARHGAP39 binds to Myo10 which moves along filopodial actin filaments. (6) Once transported to the tip, ARHGAP39 binds to mDia2 and inactivates Cdc42 which results in filopodia retraction (7).

## DISCUSSION

Localized activation of GTPases (Mathis, 1995) and their regulators (Cherfils and Zeghouf, 2013) are critical for the spatiotemporal regulation of cell protrusion (Machacek et al., 2009) and consequent cell motility. Using scaffold proteins to target active complexes to specific sites in cells is one strategy used by the cell to stimulate signaling proteins precisely in time and space. Our findings identify WAVE and mDia2 as scaffold proteins for ARHGAP39, a Rac and Cdc42 GAP. Activation of the WRC at the membrane has been well characterized (Lebensohn and Kirschner, 2009; Mendoza et al., 2011), but the precise mechanism of inactivation has remained unclear. WAVE disengagement from the membrane has been described as a prerequisite step for mDia2 activation and the formation of filopodia (Beli et al., 2008). Our data establish that ARHGAP39 activation upon binding to WAVE and Src phosphorylation on residues Y313,325 decreased Rac activation, thus controlling WRC inactivation and mDia2-mediated filopodia formation. This indicates the need for a GAP activity specific toward Rac in the WRC and supports a role for ARHGAP39 in controlling lamellipodia/filopodia coordination. Furthermore, ARHGAP39 binding to WAVE precedes its phosphorylation on Y313,325, thus suggesting that the interaction with WAVE changes the structural conformation of ARHGAP39 to make the two tyrosine residues accessible to Src. Further studies of the structure of the ARHGAP39-WAVE complex will be needed to examine this hypothesis.

Similar to WAVE, much is known about the activation of mDia2 through binding of active Cdc42 to the GBD region that disrupts intramolecular autoinhibition(Alberts, 2001), but a mechanism of Cdc42/mDia2 inactivation in filopodia has remained unclear. We demonstrate here that ARHGAP39 phosphorylation on residue Y785 by Src is critical for its interaction with the carrier protein Myo10 and its subsequent transport within filopodia. Once at the filopodia tip, ARHGAP39 can bind the FH1 region of active mDia2 that places ARHGAP39 in proximity to the active GTPase binding site on mDia2. We do not know the activity status of ARHGAP39 transported by Myo10. However, it is established that active Src localizes to the filopodia tip (Jacquemet et al., 2016) suggesting that ARHGAP39 could be activated by Src once ARHGAP39 interacts with mDia2. We show that accumulation of ARHGAP39 at filopodia tip precedes filopodia retraction and that decreased Cdc42 activity at filopodia tip leads to filopodia shortening, suggesting that Cdc42 must remain active and bound to mDia2 for actin polymerization to occur. Therefore, the immediate activation state of Cdc42 predicts filopodia growth or retraction. This is corroborated with the significant decrease in the number of retracting filopodia and increase in filopodia length in ARHGAP39-depleted cells. We did not observe enhanced Cdc42 activity at the filopodia tip in the absence of ARHGAP39, suggesting that Rac/WAVE is still active and the majority of mDia2 is trapped with the WRC; therefore, the amount of mDia2 available for Cdc42 could be limited. Neither the WAVE-binding deficient ARHGAP39 nor the Y313,325A non-phosphorylatable mutants are capable of generating filopodia, implying that the GAP function of ARHGAP39 may be sequential: first as a Rac GAP in the active WRC, then as a Cdc42 GAP in the active mDia2 complex. Our data reconcile two sets of findings: activated mDia2 is necessary for filopodia extension and its activation depends on binding to active Cdc42 (Schirenbeck et al., 2005), yet no Cdc42 activation has been detectable in filopodia (Nalbant et al., 2004). Our results provide cellular confirmation of Cdc42 localization at filopodia tips as seen in filopodia-like structures (Lee et al., 2010) and reveal the Cdc42 activation state in dynamic filopodia. We also observe discrete areas of active Cdc42 on the plasma membrane that appear to mark the area of filopodia growth immediately prior to filament extension. Future studies will be needed to determine if these foci are enriched with toca-1 and activated N-WASP prior to mDia2 activation as predicted by the clustering outgrowth model (Lee et al., 2010).

Rho family GTPases are essential in the regulation of cell behavior and cell morphology, and the deregulation of their expression or activity is associated with many human tumor types (Sahai and Marshall, 2002). Because GAPs inhibit GTPase activity, it is legitimate that GAPs that are frequently inactivated or downregulated in human cancers are considered as tumor suppressors (Vigil et al., 2010) as DLC1 (deleted in liver cancer, known also as ARHGAP7) (Xue et al., 2008; Yuan et al., 1998) or GRAF (GTPase regulator associated with focal adhesion kinase-1, known also as ARHGAP26) (Borkhardt et al., 2000; Hildebrand et al., 1996; Yao et al., 2015). In contrast, ARHGAP39 gene is upregulated in most human cancers in the Cancer Genome Atlas (TCGA) database, its high expression level is associated with poor survival (http://www.oncolnc.org) and might be considered as a potential marker for gastric cancer (Li et al., 2021). Depletion of ARHGAP39 in cells or expression of GAP inactive ARHGAP39 significantly reduces cell invasion in vitro and in mice. Overall, our data suggest that, by controlling lamellipodia/filopodia synchronization and filopodia dynamics, ARHGAP39 may be a tumor promoter gene for multiple cancer types.

## ACKNOWLEDGMENTS

We would like to thank R. Cheney for providing the Myo10 construct, A. Pendergast for providing the GFP-mDia2 and ΔDAD-mDia2 constructs, M. Innocenti for providing the WAVE2 construct, K. Burridge for providing the Vav2 DH/PH domain construct, L. Hodgson for providing the Rac1 FRET probe, and B. Felding for providing the MDA-MB435 cell lines stably expressing Firefly luciferase. We thank Rianna Flores for help with the 3D cell migration and intercalation assays and Mary MacGuckian for help with the IVIS. M.H. was supported by the training grant number T32AI007244 from NIH/NIAID. We thank NIH for funding to K.M.H (NIH R35GM122596) and to C.D.M. (NIH GM099837).

## AUTHOR CONTRIBUTIONS

M.H. conceived the initial identification of ARHGAP39 and screen of the WW domain pull-down by mass spectrometry. M.H. and C.D.M. conceived and designed the full study and wrote the manuscript. M.H. and V.D.W. carried out the experiments. V.D.W. carried out the mice experiments. M.H. and V.D.W. wrote the methods and the figure legends. C.W. and K.M.H. designed the Cdc42 FRET probe. C.W. carried out the FRET experiments and wrote the FRET methods. R.P. carried out the RBD and PBD experiments. M.H., VD.W. and C.D.M. analyzed all the data. M.H., V.D.W., C.W., K.M.H. and C.D.M. analyzed the FRET data. C.D.M. supervised the study. M.H., V.D.W., K.M.H. and C.D.M. edited the manuscript. All authors approved the final version of the manuscript.

## STAR METHODS

### RESOURCE AVAILABILITY

#### Lead contact

Further information and requests for resources and reagents should be directed to and will be fulfilled by the Lead Contact, Céline DerMardirossian.

#### Materials availability

All materials generated in this study are available on request to Lead Contact.

#### Data and code availability

This study did not generate/analyze datasets or codes.

### EXPERIMENTAL MODEL AND SUBJECT DETAILS

#### Cell culture

HeLa, HT-1080, EA.hy926 and MDA-MB-435 cells were cultured in Dulbecco’s modified Eagle’s medium (DMEM, Gibco) containing 25 mM HEPES (Gibco), 8% Fetal Bovine Serum (Gemini Bio-Products, CA), 2 mM L-glutamine (Corning) and 50 μg/ml gentamycin (Quality Biological^TM^) at 37°C in presence of 5% CO_2_. MDA-MB-231 cells were grown in the same medium supplemented with non-essential amino-acids. MDA-MB-435 were obtained from Dr. Felding (The Scripps Research Institute) and are stably transduced with Firefly luciferase to analyze tumor growth and metastasis by non-invasive bioluminescence imaging (Santidrian et al., 2013). For Cdc42 FRET experiments, HeLa cells and 293T cells were obtained from the Lineberger Comprehensive Cancer Center tissue culture facility (UNC Chapel Hill) and were cultured in DMEM (high glucose, with glutamine, Mediatech) supplemented with 10 % fetal bovine serum (HyClone) and antibiotics (Mediatech).

#### In vivo animal studies

Female 6- to 8-week old nude mice (Jackson Laboratory) were injected with 5×10^5^ F-luciferin-tagged MDA-MB-435 control or Wdom stable tumor cells into the tail vein for experimental metastasis or subcutaneously into the fourth axillary mammary fat pad for primary tumor growth. Mice were imaged weekly (IVIS 200; Xenogen) 10 min after intraperitoneal injection of D-luciferin (150 mg/kg).

Bioluminescence was quantified as photons/s/cm^2^/sr in defined regions of interest using Living Image software. For primary tumor growth, tumor size was also monitored every week externally using a caliper. Mice were euthanized after 12 weeks, mammary pad tumors were removed surgically and weighed. For experimental metastasis, mice were euthanized after 15 weeks.

*Study approval*. Animal studies were approved by the Institutional Animal Care and Use Committee of The Scripps Research Institute. Animal work complied with NIH and institutional guidelines (The Scripps Research Institute is AAALAC accredited).

### METHOD DETAILS

#### Plasmids

Myo10 was a gift from Dr. Cheney, GFP-mDia2 full-length and deltaDAD from Dr. Pendergast and WAVE2 from Dr. Innocenti. The Vav2 DH/PH domain plus C-terminal regulatory domain (amino acids 184-878) was a gift from Dr. Burridge. Full-length ARHGAP39 was cloned into pCMV5-HA3, pEGFP-C2 and pTriEx-4 Flag-mCherry. Nterm domain of ARHGAP39 (aa 1-134) was amplified by PCR and cloned into pCMV5-HA3 and pGEX4T1. A shorter Nterm fragment (aa 23-97) was amplified by PCR, cloned into pBabe puro with and without GFP and stably transfected into tumor cells. HA-ARHGAP39 mutants were amplified by PCR: Y313A; Y325A; Y785A; Y313, 325A (2YA); Y313, 325, 785A (3YA); Wmut (W53A, P56A, W92A, P95A); RA (R932A), Cterm (aa135-1084). For pTriEx-4 Flag-mCherry, full-length ARHGAP39 was amplified by PCR and inserted into pTriEx-4 Flag via HindIII restriction site. mCherry fluorescent protein was amplified by PCR and ligated into this vector via BamHI site upstream of HindIII. Myo10 Myth-FERM domain (aa1486-2058) was amplified by PCR and cloned into pGEX4T1. WAVE2 full-length, -Nterm (aa1-299), -Cterm (aa420-498) and -poly-proline region (PP, aa300-419) and mDia2 full-length, -deltaDAD (aa1-1057), -Nterm (aa1-514), -Cterm (aa689-1193), -FH1 domain (aa506-696) used in in vitro translation system were cloned into pCDNA3.1+ with an HA-tag sequence (ATGTACCCATACGATGTTCCAGATTACG-CT) inserted upstream of the KpnI site. Refer to Table 1 for cloning primers. All constructs were confirmed by sequencing.

**Table 1.**
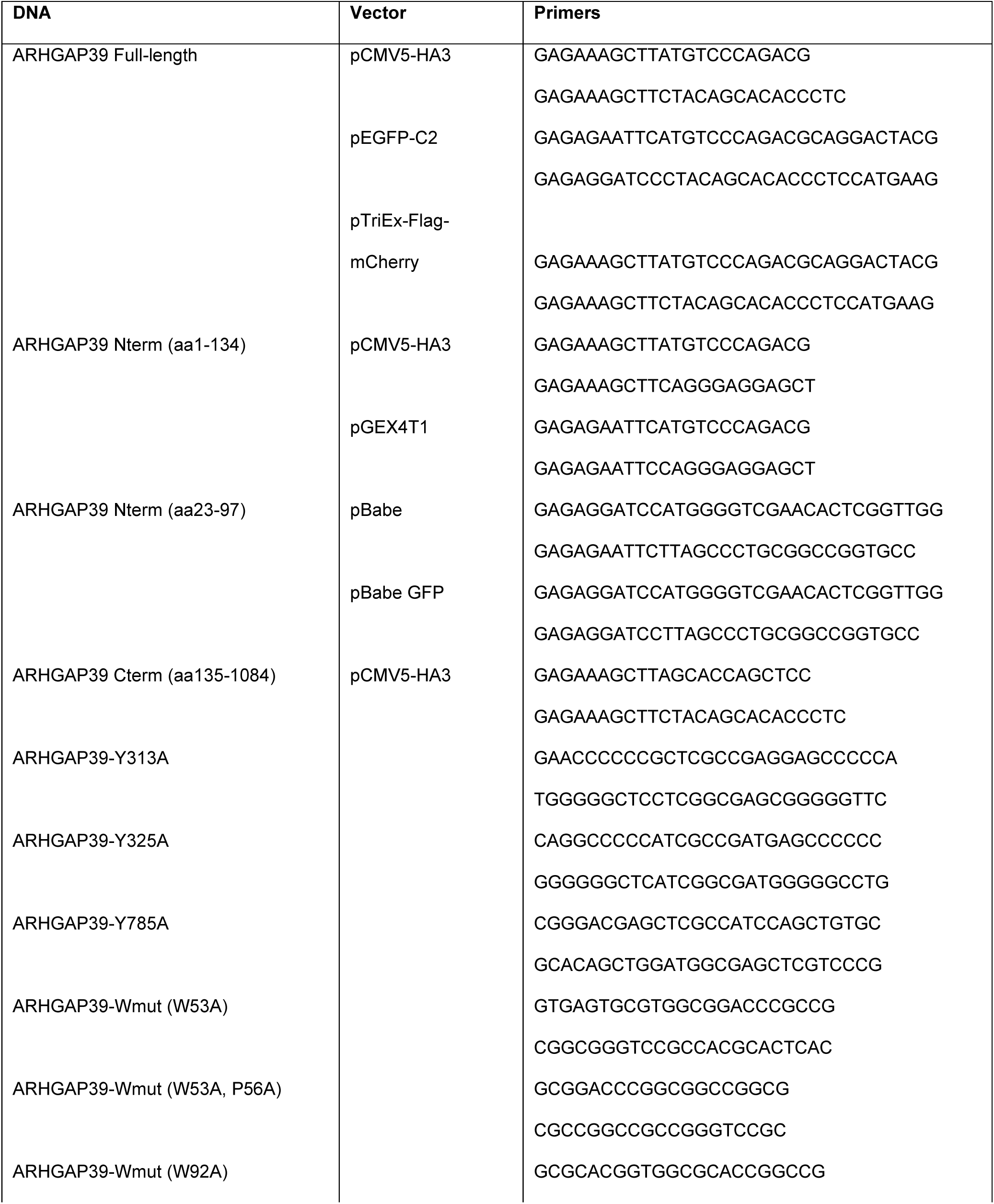
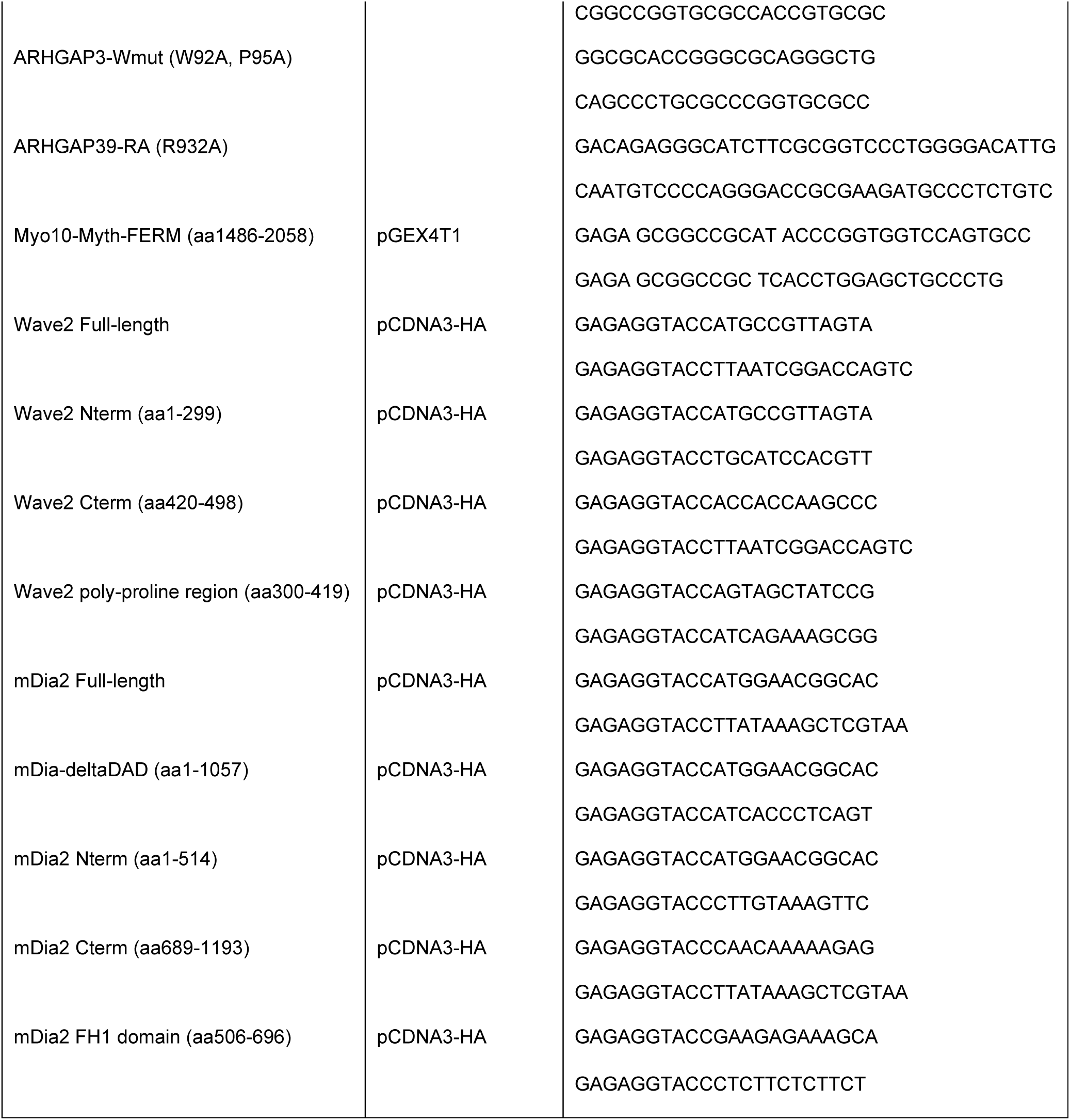
List of cloning primers.

#### RNA interference

Synthetic siRNA oligos for ARHGAP39 (5’-CAAGGUCGCUGAUCGGGAGUU-3’), mDia2 (5’-GCUCAAACCUUCGGAUUUAUU-3’) and nontargeting control oligos (5’-GCGCGCUAUGUAGGAUUCGUU-3’) were purchased from Dharmacon. siRNAs were transfected into cells at 20 nM concentration with Lipofectamine 2000 (Invitrogen) for 48 to 72 h. For rescue experiments, silent mutations (V217 GTC→GTA and R220 CGG→CGA, noted ARHGAP39 siR) were introduced into siRNA-targeted region of ARHGAP39 by PCR.

#### Antibodies and reagents

The following antibodies were used: mouse monoclonal anti-HA.11 (BioLegend), rabbit monoclonal anti-WAVE2 (D2C8, Cell Signaling), mouse monoclonal anti-GFP (3E6, Molecular Probes), mouse monoclonal anti-Rac1 (23A8, Upstate Biotechnology), rabbit polyclonal anti-Cdc42 (Cell Signaling), mouse monoclonal anti-RhoA (Santa Cruz Biotechnology Inc.), rabbit polyclonal anti-c-src (SRC2, Santa Cruz), rabbit monoclonal anti-phosphoSrc (Y416, Millipore), mouse monoclonal anti-actin C4 (Millipore). Goat anti-mouse or anti-rabbit Alexa Fluor-488 or −647 conjugated secondary antibodies, Alexa Fluor-350 and −633 conjugated phalloidin were obtained from Molecular Probes. Human recombinant TNFα was obtained from R&D Systems and was used at a final concentration of 20 ng/ml.

##### Cell transfection

(Santidrian et al., 2013)Transient and stable DNA transfections were performed using Lipofectamine 2000 (Invitrogen), according to the manufacturer’s instructions. Stable cell lines were selected with 1 μg/ml puromycin (Invitrogen). For Cdc42 FRET experiments, DNA plasmids were transfected into HeLa cells using Fugene6 according to the manufacturer’s instructions. To test for ARHGAP39 siRNA efficiency, cells transfected with 20 nM siRNA oligos control or ARHGAP39 for 48 h were transfected again with siRNA oligos and HA-ARHGAP39 -WT or –siRNA resistant (-siR). 24 h later, cells were lysed in 25 mM TrisCl pH 7.5, 1 mM EDTA, 1 mM MgCl2, 150 mM NaCl, 1% NP-40 in presence of protease inhibitors. HA-ARHGAP39 expression in lysates was detected by immunoblotting using anti-HA antibodies.

#### GTPase activity assay

Recombinant GST-PBD and GST-RBD were purified as previously described (Huang et al., 2013). Briefly, GST-PBD and -RBD proteins were expressed from pGEX vectors in BL21, induced for 5 h at 37°C or 30°C, respectively, and lysed in native buffer (50 mM TrisCl pH 7.5, 100 mM NaCl, 5 mM MgCl_2_, 1 mM EDTA, 1mM DTT). GST proteins were bound to glutathione-sepharose, washed several times in buffer A (20 mM Hepes pH 8, 0.5 M NaCl) and eluted in 30 mM reduced glutathione in 0.3 M Hepes pH 8.0. Recombinant proteins were dialyzed in 25 mM TrisCl pH 7.5, 1 mM EDTA, 5 mM MgCl2, 50 mM NaCl, 5% Glycerol, 0.5 mM DTT. Rac-, Cdc42- and RhoA-GTP levels were measured by PBD and RBD pulldowns, respectively, as previously described (Pathak et al., 2012; Stofega et al., 2006). HeLa cells transfected with control or ARHGAP39 siRNA oligos for 48 h were lysed in lysis/assay buffer (20 mM Hepes pH 7.5, 150 mM NaCl, 10 mM MgCl_2_, 5% Glycerol) and a portion of the lysates were retained for immunoblotting. Lysates were incubated with 15 μg of recombinant purified GST-PBD or -RBD and glutathione sepharose for 1 h at 4°C, washed in lysis/assay buffer and boiled in SDS-sample buffer. Co-precipitated Rac, Cdc42 or RhoA were visualized by immunoblotting and the amount of active GTPase was normalized to the total amount in cell lysates.

#### In vitro translation and binding assays

To determine whether ARHGAP39 directly binds to WAVE and mDia2, recombinant GST and GST-Wdom were expressed and purified as described above for recombinant GST proteins. HA-WAVE and HA-mDia2 constructs were in vitro translated using TNT Quick Coupled Transcription/Translation system (Promega). Briefly, vectors encoding the proteins of interest were incubated with the TNT Quick Master Mix for 90 min at 30°C in a 50 μl reaction volume. 12 μl of the expressed proteins were used in subsequent binding reactions. Extracts were incubated with GST or GST-Wdom in binding buffer (50 mM TrisCl pH 7.4, 150 mM NaCl, 1% NP40) and glutathione sepharose for 2 h at 4°C. Samples were washed three times in binding buffer, resolved by SDS-PAGE and immunoblotted with anti-HA antibodies. To determine whether ARHGAP39 binds to Myo10, recombinant GST-Myo10 Myth-FERM domain was expressed from pGEX vectors in BL21, induced with 0.3 mM IPTG for 3 h at 37°C and lysed in lysis buffer (50 mM TrisCl pH 7.5, 1 mM EDTA, 5 mM MgCl2, 150 mM NaCl) containing 1 mM DTT, 1 mg/ml lysozyme, 20 U/ml DNAseI. Lysates were sonicated 5x 30 s on ice and clarified by centrifugation. GST proteins were bound to gluthatione-sepharose and washed several times in lysis buffer. HeLa cells transfected with HA-ARHGAP39 -WT, −785A or −2YA in combination with active Src (Src YF) were lysed in 25 mM TrisCl pH 7.5, 1 mM EDTA, 1 mM MgCl2, 150 mM NaCl, 1% NP-40 in the presence of protease inhibitors and sodium orthovanadate. A portion of the lysates were retained for immunoblotting. Lysates were incubated with 20 μg of recombinant purified GST or GST-Myo10 Myth-FERM on gluthatione-sepharose for 1 h at 4°C, washed in lysis buffer and boiled in SDS-sample buffer. Co-precipitated ARHGAP39 was visualized by immunoblotting using anti-HA antibodies.

#### Phosphorylation assay

HeLa cells were transfected with ARHGAP39-WT or various mutants in combination with empty vector, active Src (Src YF) or inactive Src (Src KM). 16 h post-transfection, cell extracts were prepared in lysis buffer containing 25 mM TrisCl pH 7.5, 5 mM MgCl_2_, 150 mM NaCl, 1% NP40, 1 mM phenylmethylsulfonyl fluoride (PMSF), and protease inhibitors (pepstatin, leupeptin and aprotinin). Mouse monoclonal anti-HA antibody was used for immunoprecipitation and protein G sepharose beads (GE Healthcare, Uppsala, Sweden) were then added. Immunoprecipitates (IPs) were washed four times in lysis buffer and eluted in SDS-sample buffer. IPs and whole cell extracts were then resolved by SDS-PAGE and analyzed by immunoblotting.

#### Immunofluorescence microscopy

16 h post-transfection with various constructs, cells were replated on fibronectin- (10 μg/ml, Sigma) or collagen- (type I, rat tail, 50 μg/ml, Upstate) coated glass coverslips and allowed to adhere for 3 h. Cells were then fixed in cytoskeletal buffer (CB; 10 mM MES, 3 mM MgCl_2_, 138 mM KCl, 2 mM EGTA, pH 6.9) containing 4% paraformaldehyde (PFA), permeabilized in CB containing 0.5% Triton X-100 and blocked with 2% BSA in CB. Cells were then immunolabeled for HA, GFP or WAVE by using the appropriate Alexa Fluor-conjugated secondary antibodies. F-actin was detected by using Alexa Fluor-conjugated phalloidin. Cells were mounted on slides with Prolong Gold antifade reagent (Molecular Probes). Epifluorescence images of fixed cells were acquired on an inverted microscope (Eclipse TE 2000-U, Nikon) equipped with an electronically controlled shutter, filter wheels, and a 14-bit cooled CCD camera (Cool SNAP HQ, Photometrics) controlled by Metamorph imaging software (Molecular Devices Inc.) by using a 60x/1.49 NA TIRF objective lens (Nikon).

#### Immunofluorescence analysis

Quantification of the fluorescence intensity of WAVE and HA-ARHGAP39 as a function of the distance from the cell edge was obtained with custom software written in Matlab (Mathworks). Bands of constant distance to the cell edge were constructed, individual fluorescence intensities were accumulated and averaged in each band to produce fluorescence intensities versus distance-to-the-cell-edge graphs. The data shown represent one experiment and are averaged for over 20 cells for each condition.

Quantification of fluorescence intensity of ARHGAP39, F-actin, Myo10 and mDia2ΔDAD inside filopodia was measured using the linescan (3 μm) function in Metamorph.

#### Cell proliferation assay

10^4^ control or ARHGAP39 Wdom-expressing stable MDA-MB-435 cells were seeded in 24-well plate, in triplicate for each time point. Every 24 h, cells were trypsinized, resuspended in 1 ml of complete culture medium and counted using an automated cell counter (TC20 automated cell counter, Bio-Rad).

#### Soft agar colony formation assay

Soft agar colony formation assay was carried out in 6-well plates. Briefly, 1% and 0.6% agar solutions were prepared, sterilized, and maintained at 40°C throughout the experiment. The bottom layer of agar was prepared by mixing 0.75 ml of 1% agar with 0.75 ml of 2x DMEM growth medium supplemented with 20% FBS and allowed to solidify at room temperature for 30 min. For the top layer, 2×10^4^ control or ARHGAP39 Wdom-expressing stable MDA-MB-435 cells were suspended in 0.75 ml of 2x DMEM medium supplemented with 20 % FBS and mixed with 0.75 ml of 0.6% agar. The top layers were allowed to solidify for 30 min at 37°C. The plates were incubated in a 5% CO2 incubator at 37°C for 3 weeks. 1 ml of DMEM growth medium was added twice a week. After 21 days, cells were imaged on an inverted microscope (Eclipse TE 2000-U, Nikon) using a 10x/0.25 Ph1 DL Phase Contrast objective lens (Nikon). Cells were then stained with 0.05% Crystal Violet for 20 min before being washed in distilled water. The number of colonies was counted for each condition.

#### 3D motility assay

24 h after being transfected with siRNA control and ARHGAP39 oligos, HT-1080 cells were trypsinized and 2×10^4^ cells were resuspended in 40 μl of complete medium. Cells were mixed into 360 μl of collagen gel solution (2 mg/ml Collagen I, 1x MEM, 2.35 mg/ml NaHCO_2_, 20 mM Hepes, 24 mM Glutamine), rapidly placed into a 4-well glass-bottom chamber (LabTek) and incubated at 37°C for 3 h. Once the collagen gels have polymerized, 500 μl of complete medium was added on top of the gels and cells were incubated overnight at 37°C. The next day, chambers were placed onto a temperature-controlled stage and phase-contrast time series were acquired on our inverted microscope using a 10x/0.25 Ph1 DL Phase Contrast objective lens. Images were captured every minute for 2.5 h. Quantification of motility parameters, including net path length, the net distance that the cells traversed from the first to the last frame; total path length, the total distance traversed by cells over time and cell velocity was performed using the Track Object tool in Metamorph.

#### Trans-endothelial migration assay

Undersides of transwells (8 μm pores, #3422, Costar) were coated with 5 μg/ml matrigel (Corning) for 1 h. 10^5^ EA.hy926 endothelial cells were then plated on top of the matrigel for 48 h at 37°C to form a monolayer. HT-1080 tumor cells transfected with empty vector control or various ARHGAP39 constructs, MDA-MB-231 or −435 stable cells expressing GFP or GFP-Wdom cells were grown overnight in serum-free medium. The following day, 2.5×10^4^ starved cells were placed into the upper chamber, in serum-free medium. Cells were allowed to migrate toward complete growth medium for 24 h at 37°C. Transwells (top and bottom) were then fixed in CB with 4% PFA and permeabilized in PBS with 0.5% Triton X-100 and 0.7% fish skin gelatin. Cells were washed and blocked in PBS containing 0.05% Triton X-100 and 0.7% fish skin gelatin. Cells were then immunostained for HA, or GFP, and F-actin. Cells were post-fixed in CB with 4% PFA before the transwell membranes were gently cut with a scalpel and mounted on slides with Prolong Gold antifade. A 22×22 mm #1 glass coverslip was placed on top of the membranes and allowed to dry before to be imaged. Images were acquired using a 40x/1.3 Oil objective lens (Nikon) on a spinning disk (Yokogawa; PerkinElmer) confocal microscope (TE 2000-U, Nikon; (Adams et al., 2003)). Z series images were acquired every 0.3 μm. Confocal z projections of the entire transwell were obtained with ImageJ 3D Volume viewer. The percentage of migrated cells was quantified by counting the number of tumor cells that have crossed the membrane and the endothelial cell layer versus the number of cells that have remained on top of the transwell.

#### Intercalation assay

3×10^5^ EA.hy926 endothelial cells were plated on 15 mm fibronectin-coated glass coverslips (10 μg/ml) for 48 h at 37°C to form a monolayer. 7×10^4^ MDA-MB-231, −435 or HT-1080 tumor cells, transiently or stably transfected, were added on top of EA.hy926 monolayer. Tumor cells were allowed to interact with endothelial cells for 180 min before being fixed and labelled for HA, or GFP, and F-actin. Z-stack images were acquired on a spinning disk confocal microscope as described above. Confocal z projections of cells were obtained with ImageJ 3D Volume viewer. The percentage of intercalated cells was quantified by counting the number of tumor cells that have integrated into the endothelial cell monolayer versus the number of cells that have remained on top of the endothelial cells.

#### Live cell imaging

For ARHGAP39 dynamics in filopodia, HT-1080 cells were transfected with GFP-ARHGAP39 in combination with myc-tagged Cdc42-WT. 16 h post-transfection, cells were replated on collagen-coated glass coverslips and allowed to adhere for 3 h. Coverslips were then mounted into a chamber with phenol red free medium (Gibco) supplemented with 10% FBS and were placed onto a temperature controlled stage during observation. GFP time series were acquired using a 60X/1.49 NA TIRF objective lens (Nikon) on a spinning disk confocal microscope (TE 2000-U, Nikon). Images were acquired every 5 s for 10 min. For filopodia dynamics in siRNA ARHGAP39 cells, HeLa cells were transfected with siRNA control or ARHGAP39 oligos for 24 h before being transfected with GFP-Myo10, to induce the formation of filopodia, in combination with siRNA oligos. 16 h post-second transfection, cells were replated onto fibronectin-coated glass coverslips, allowed to adhere for 3 h and mounted on chambers as described above. Phase-contrast time series were acquired using a 100X/1.4 NA Plan APO objective lens (Nikon) on a spinning disk confocal microscope (TE2000-U, Nikon). Images were acquired every 3 s for 15 min. Filopodia lifetime was calculated for filopodia that form and then totally disappear during the timeframe of the movie, whether they retract, collapse or get embedded by the protruding membrane. Filopodia length was measured using Metamorph imaging software.

#### FRET imaging

##### Rac1 FRET

The single chain, optimized FRET Rac1 biosensor was kindly provided by Dr. Hodgson (Albert Einstein College of Medicine, Bronx, NY)(Moshfegh et al., 2014). HeLa cells were transfected with the Rac1 biosensor in combination with empty vector control or various HA-tagged ARHGAP39 constructs. 16 h post-transfection, cells were replated on fibronectin-coated glass coverslips and allowed to adhere for 3 h before being fixed in CB with 4 % PFA. Cells were then immunolabeled for HA tag and mounted on slides with Prolong Gold antifade. Images were acquired using a 60X/1.49 NA TIRF objective lens (Nikon) on our inverted microscope (Eclipse TE2000-U, Nikon) using two 14-bit cooled CCD cameras (Cool SNAP HQ, Photometrics). CFP and FRET emission channels were acquired simultaneously using a dual-emission image splitter system (CAIRN Research). The following bandpass filters (Chroma Technology) were used: ET436/20X for CFP excitation, ET480/40M for CFP emission and ET535/30M for FRET emission. A 505 dclp beam splitter (Chroma Technology) was used to separate CFP and FRET emissions. FRET efficiency was calculated using Metamorph (Molecular Devices Inc.) and Matlab (Math Works Inc.) software as described in Spiering et al(Spiering et al., 2013). Briefly, CFP and FRET images were corrected for shading and dark current, then background subtracted. Images were then intensity thresholded and masked. Finally, the FRET ratio was calculated by dividing the FRET image by the CFP image using a scaling factor of 1000. Rac1 FRET efficiency at protrusions was measured from the cell edge to 3 μm inside the protrusion. FRET efficiency was normalized to Rac1 activity in control cells expressing empty vector.

##### Cdc42 FRET

###### Production of the intermolecular Cdc42 biosensor

For Cdc42 biosensor imaging, we used a dual-chain biosensor generated by expressing plasmids encoding either Cdc42 fused to the C terminus of mCerulean, a CFP variant optimized for FRET(Nguyen and Daugherty, 2005), or the Cdc42-binding CRIB domain from WASP (CBD), amino acids 230–314, fused to the C terminus of YPet, a YFP variant optimized for FRET (Extended Data Figure 4a)(Nguyen and Daugherty, 2005). This was based on our earlier biosensor designs, now using optimized fluorescent proteins for FRET. The EGFP coding region from the EGFP-C1 vector (Clontech, Inc.) was replaced with a PCR product containing the mCerulean or YPet coding regions flanked by an NcoI restriction site and a SGLRSELGS linker containing a BamHI restriction site. The PCR products of the Cdc42 and CBD coding sequences were inserted between the BamHI restriction site in the SGLRSELGS linker and an EcoRI restriction site in the downstream multiple cloning sites of the vector.

##### Fluorometry assay of Cdc42 activation

The biosensor was validated by performing FRET measurements in a spectrophotometer, using cells in suspension. In brief, 293T cells were plated on poly-L-lysine coated wells of a 6-well dish 36 h before the assay was run. Cells were transfected using Lipofectamine according to the manufacturer’s instructions. A total of 650 ng of DNA was transfected per well, consisting of 100 ng of mCerulean-Cdc42, 100 ng of YPet-CBD, and the balance consisted of Vav2 DH-PH domain or Rho-GDI-1 (used as positive and negative controls for Cdc42, respectively) or blank vector DNA. Each experimental condition was set up in triplicate. 24 h post-transfection, cells were checked for appropriate brightness and cell health, washed once with DPBS, trypsinized, and then resuspended in 500 μL cold DPBS. 400 μL of cell suspension was then loaded into a spectrophotometer cuvette. Using a SPEX Fluorolog spectrophotometer, the cell suspension was excited at 433 nm, the excitation of mCerulean, and the YPet emission was monitored from 450 nm to 600 nm at 3 nm intervals. For each test, a sample transfected with YPet-CBD only was monitored to account for bleed-through into the FRET channel from direct excitation of the acceptor fluorophore. This reading was subtracted from each measurement. For the calculation of FRET ratio, after bleed-through correction, each sample was normalized such that the peak of CFP emission (475 nm) was set as an intensity value of 1.0. The FRET emission peak (525 nm) was then divided by the CFP peak for each sample, and the samples were averaged for each condition. Statistical significance between conditions was assessed by two-tailed Student’s *t*-test, corrected by Bonferroni’s principle. The biosensor was validated by measuring the FRET efficiency in cell lysates expressing CBD-YPet and constitutively active ECFP-Cdc42 (QL) or wild-type ECFP-Cdc42 (WT), in the presence of either empty vector YPet, Vav2, a Cdc42 GEF activator(Abe et al., 2000) or GDI, a GTPase inhibitor(DerMardirossian and Bokoch, 2005) (Extended Data Figure 4b). FRET normalized to Cdc42-WT probe alone increased in cells expressing active Cdc42 or Vav2 but decreased in cells overexpressing GDI.

##### Induction of filopodia during imaging

To study the activation of Cdc42 in filopodia, cells were transfected with control or ARHGAP39 siRNA oligos 48 h prior to imaging, and then subsequently transfected with the Cdc42 biosensor 36 h prior to imaging (Gadea et al., 2004). Cells were then plated on fibronectin-coated coverslips 24 h prior to imaging. Six hours before imaging, cells were starved in Ham’s F12K medium containing 0.5% delipidated BSA plus glutamine. Cells were transferred to a heated open chamber apparatus to allow for addition of 20 ng/mL TNFα. Cells were imaged for 5 min at 20 s intervals prior to the addition of TNFα and imaged for a subsequent 15 min post-stimulation to observe filopodia formation and retraction.

##### Imaging of Cdc42 activity in cells

For emission ratio imaging, the following filter sets were used (Chroma): ECFP: D436/20, D470/40; FRET: D436/20, HQ535/30; EYFP: HQ500/20, HQ535/30. A dichroic mirror (“Quad-Custom” Lot# 511112038) was custom manufactured by Chroma Technology Corp. for compatibility with all these filter sets. Cells were illuminated with a 100 W Hg arc lamp through a 10% transmittance neutral density filter. At each time point, three images were recorded with the following exposure times: mCerulean (1.2 s), FRET (excitation of donor, observation of acceptor emission) (1.2 s), YPet (0.4 s) at binning 2×2 using a 40x objective. Ratio calculations to generate activity images were performed following bleed-through correction methods described previously(Kraynov et al., 2000). Briefly, Metamorph software was used for image alignment and ratiometric calculation of activation signals. All images were shading-corrected and background-subtracted. Binary masks with values equal to 1 inside the cell and 0 elsewhere were extracted by applying a threshold to the mCerulean image, because it had the largest signal-to-noise ratio. Control cells expressing either mCerulean alone or YPet alone were used to obtain bleed-through coefficients α and *β* in the following equation:

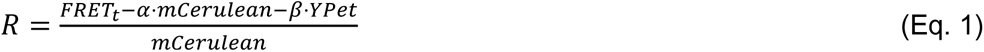

where *R* is the Ratio, *FRET_t_* is the total FRET intensity as measured, *α* is the bleed-through of minto FRET channel, *β* is the bleed-through of YPet into the FRET channel and mCerulean is the total mCerulean intensity as measured. By calculating the linear slope of the relationship between FRET and mCerulean intensities upon mCerulean excitation of cells expressing only the mCerulean, the bleed-through parameter *α* was determined. Similarly, by determining bleed-through into the FRET channel upon mCerulean excitation of cells expressing only the YPet, the bleed-through contribution of YPet excitation by mCerulean into the FRET channel *β* was determined. The *α* parameter was found to be within 0.4-0.5 and the *β* parameter was typically ∼0.2. Both were dependent on the particular optical configuration of the microscope used. With these parameters, the ratio of corrected FRET over mCerulean was calculated and used as a measure of Cdc42 activation. In time-lapse experiments, mCerulean and YPet bleached at different rates, so the ratio was corrected for photobleaching as described elsewhere (Hodgson et al., 2005). Briefly, by algebraic manipulation, (Eq.1) can be rearranged to:

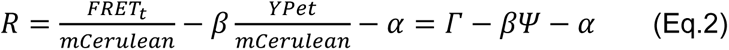

where Γ is the fraction of FRET intensity over mCerulean intensity, Ψ is the fraction of YPet intensity over mCerulean intensity, and *α* and *β* are the bleed-through constants described above. By taking double exponential fits of the decays of both Γ and Ψ, the correction function, 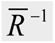, was calculated following the methods outlined in (Hodgson et al., 2005).

For visual representations of ratiometric images, a linear pseudocolor lookup table was applied to all ratio images and the ratio values were normalized to the lower scale value. In each experiment, CFP, YFP, and FRET images were carefully inspected to verify that all portions used to create the ratio image had a sufficiently high signal/noise ratio. We targeted at least 400 gray level values (12 bit dynamic range) above background in the lowest intensity regions within the cell. This was especially important in thin parts of the cell where fluorescence was low.

##### Cdc42 FRET analysis in filopodia

Filopodia in each state were isolated by applying the region drawing tool in Metamorph to the DIC image, and the regions were used to mask the FRET ratio image to quantify filopodial activity within that region. The first 8 pixels/2.6 μm were used as a cut-off to avoid error from differing filopodia lengths. For quantification of the positioning of the most common maxima of Cdc42 signal along the filopodia, 3 pixel wide linescans were performed along the long axis of filopodia on the FRET ratio image and on the CFP fluorophore control. The location of local FRET maxima along the line was determined. For quantification of Cdc42 activity in forming filopodia, at membrane regions where single filopodia formed, FRET ratio intensity in a region incorporating the base of the filopodia was measured one frame prior to filopodia formation and compared with FRET ratio intensity in a nearby region of plasma membrane of similar size.

## QUANTIFICATION AND STATISTICAL ANALYSIS

Statistical analysis was performed using two-tailed unpaired Student’s *t*-test and differences with p < 0.05 were considered statistically significant. Data are presented as mean ± SEM of values obtained in independent experiments.

## Supplementary figure legends

**Figure S1. Related to Figure 1.**
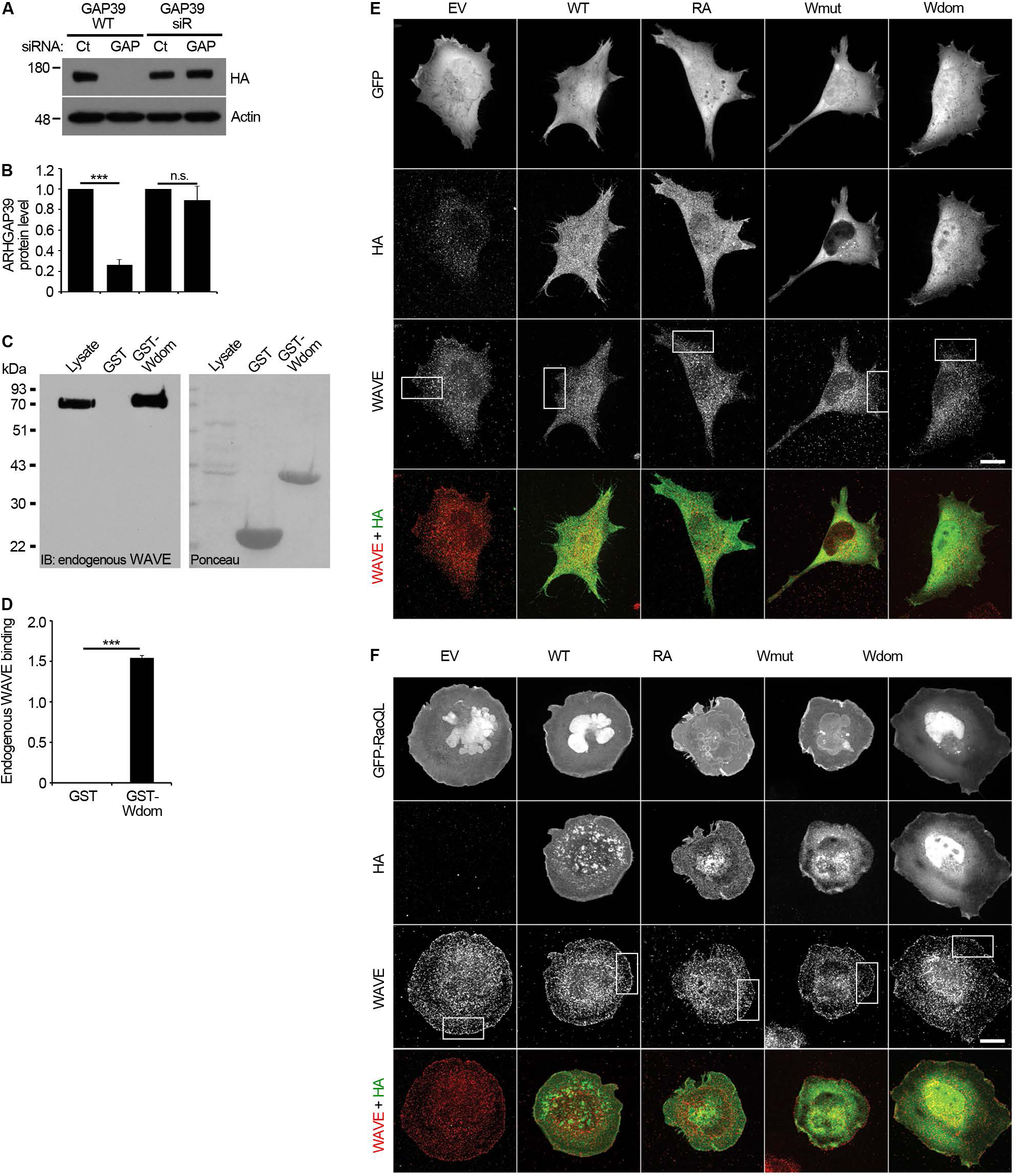
ARHAGP39 binds to endogenous WAVE and is recruited at the cell edge by active Rac. (A and B) Depletion of ARHGAP39 using siRNA oligos. (A) HeLa cells were transfected with 20 nM of either control siRNA or ARHGAP39-specific siRNA oligos. After 48 h, cells were co-transfected with 20 nM siRNA oligos and HA-ARHGAP39 -WT or -siRNA resistant (siR). Equal amounts of cell lysates were loaded and analyzed by western blot. ARHGAP39 was detected using anti-HA antibodies. Actin was used as loading control. (B) Quantification of ARHGAP39 protein levels normalized to actin levels and expressed relative to that of control siRNA. (C and D) ARHGAP39 N-terminal domain interacts with endogenous WAVE. (C) HeLa cell lysates were incubated with GST or GST-ARHGAP39 aa1-134 (noted GST-Wdom) and WAVE binding was detected using an anti-WAVE antibody. Data are representative of one experiment out of three independent experiments with similar results. (D) Quantification of endogenous WAVE binding. ***p < 10^−6^ compared to GST. (E and F) ARHGAP39 is recruited at the cell leading edge by active Rac. HeLa cells were transfected with GFP (E) or GFP-RacQ61L (active Rac) (F) in combination with either HA empty vector (EV), HA-ARHGAP39 WT, HA-ARHGAP39 RA (GAP inactive), HA-ARHGAP39 Wmut (WAVE-binding deficient) or HA-Wdom (ARHGAP39 aa 1-134). Cells were fixed and immunostained with anti-HA and anti-WAVE antibodies to visualize ARHGAP39 and endogenous WAVE, respectively. Scale bars, 15 μm. White boxes in the WAVE images indicate regions selected for magnified images shown in Figure1. Bottom panels show overlay of HA (green) and WAVE (red). Data are representative of three independent experiments. Data are presented as mean ± SEM. Statistical analysis was performed using two-tailed Student’s *t*-test. n.s., not statistically significant; *\**p < 0.05; *\*\**p < 0.01 and \**\*\**p < 0.001 unless specified.

**Figure S2. Related to Figure 2.**
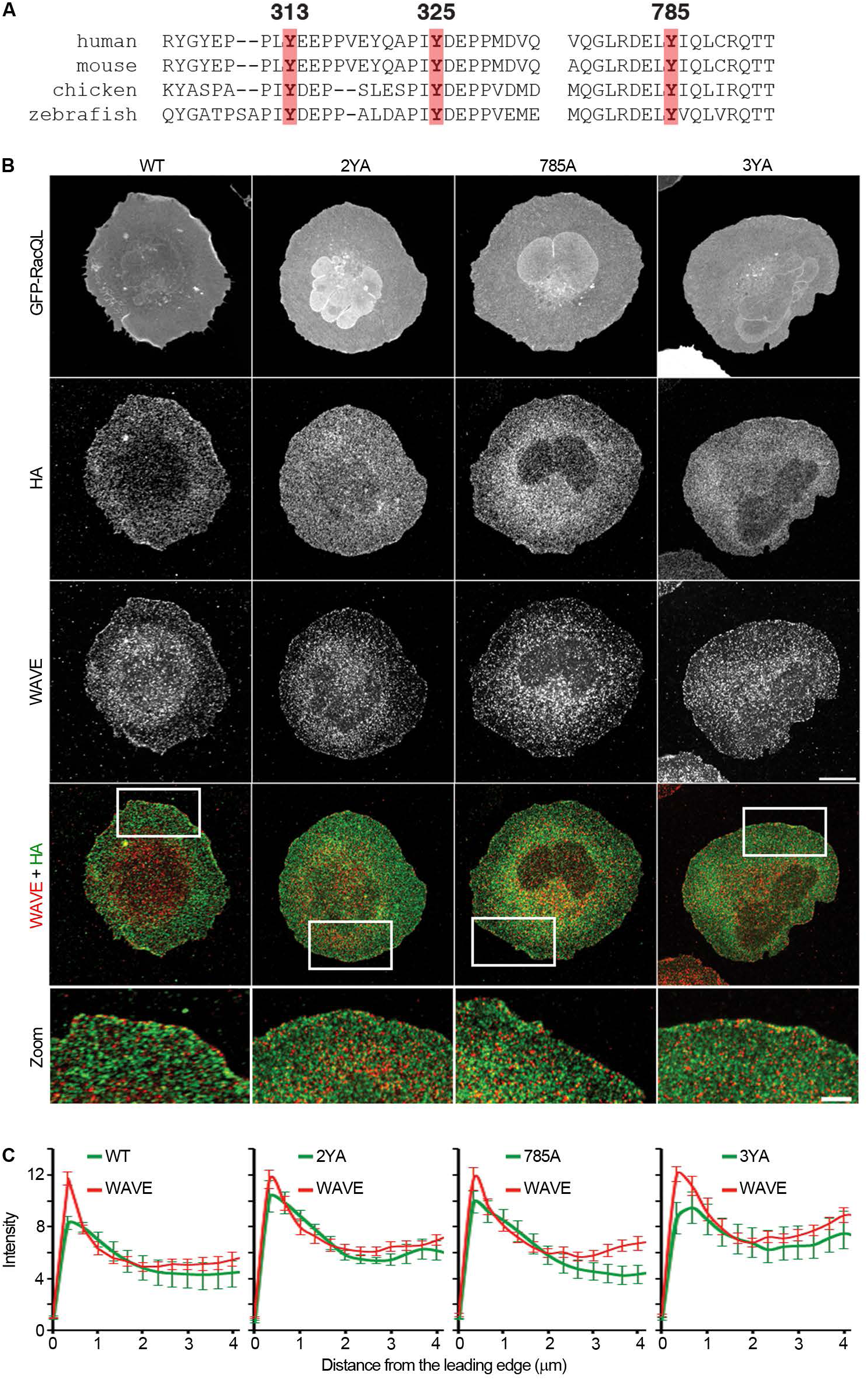
Non-Phosphorylatable mutants of ARHGAP39 are still recruited at the cell edge by active Rac. (A) Sequence alignment of ARHGAP39 in human, mouse, chicken, and zebrafish. Red boxes indicate conserved tyrosine residues. (B and C) ARHGAP39-WT and non-phosphorylatable mutants co-localize with WAVE at the cell edge. (B) HeLa cells were transfected with GFP-RacQ61L in combination with HA-ARHGAP39-WT, HA-ARHGAP39 −2YA (Y313, 325A), −785A (Y785A) or −3YA (Y313, 325, 785A). Cells were labelled with anti-HA and anti-WAVE antibodies to visualize ARHGAP39 (green) and endogenous WAVE (red), respectively. Scale bars, 15 μm. White boxes in the overlay images indicate regions selected for magnified images shown in bottom panel. Scale bars, 5 μm. (C) Average fluorescence intensity of HA (green) and WAVE (red) from cells expressing RacQL. Intensity profiles start at the cell leading edge (0 μm) and end at 4 μm inside the protrusion. n ≥ 20 cells for each condition. Data are representative of three independent experiments and are presented as mean ± SEM.

**Figure S3. Related to Figure 3.**
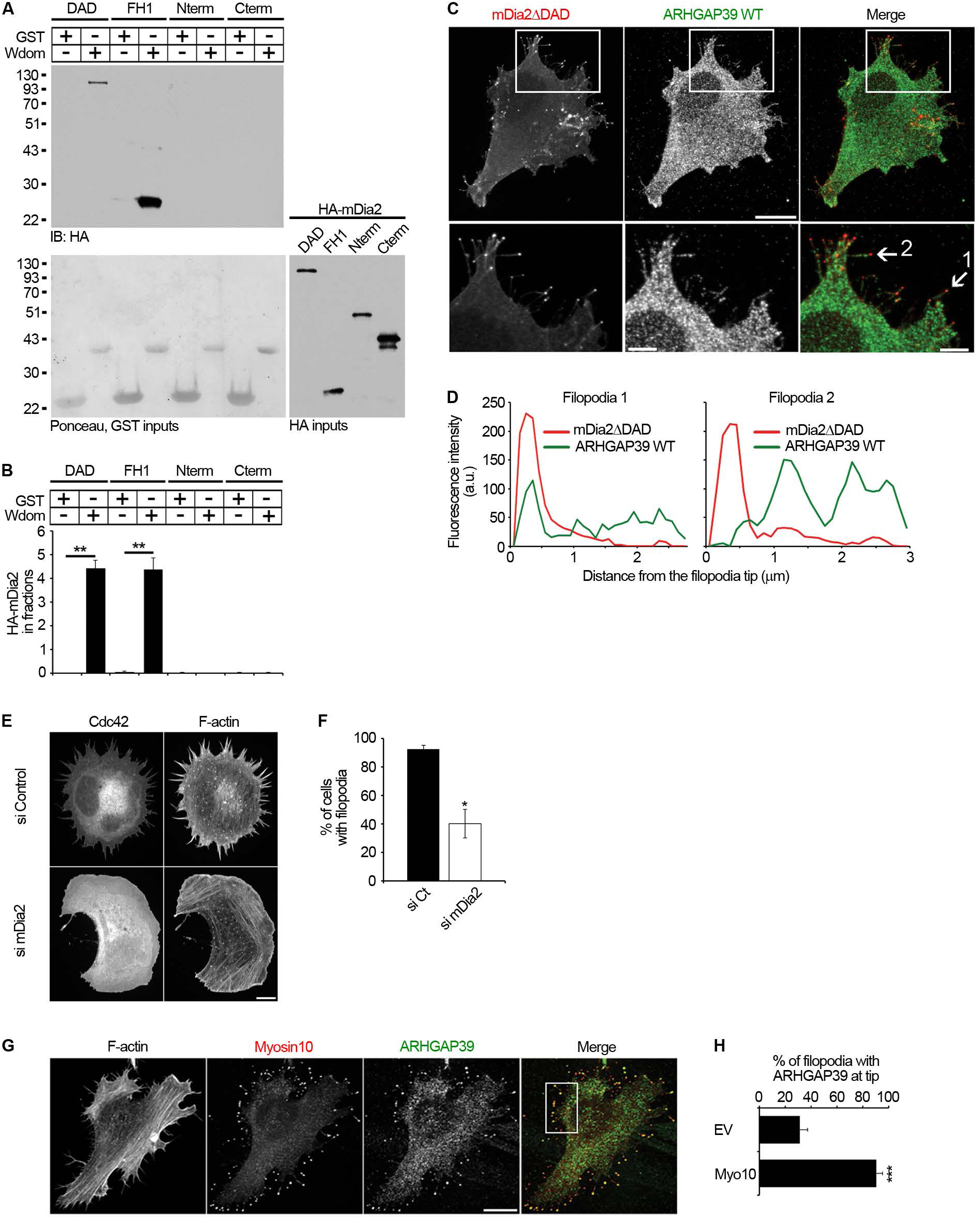
ARHGAP39 binds to mDia2 but its localization in filopodia is mDia2-independent. (A and B) ARHGAP39 N-terminal domain directly interacts with mDia2 FH1 region. (A) In vitro translated HA-mDia2 constructs deltaDAD (DAD), FH1, N-terminal (N) and C-terminal (C) were incubated with GST or GST-ARHGAP39 aa1-134 (Wdom). mDia2 proteins were detected using anti-HA antibodies. Bottom left panel, red ponceau stain of GST inputs. Bottom right panel, HA-mDia2 inputs. (B) Quantification of mDia2 binding to ARHGAP39-Wdom. (C and D) ARHGAP39 only partially co-localizes with mDia2 in filopodia tips. (C) HeLa cells were transfected with GFP-mDia2ΔDAD (active mDia2) in combination with HA-ARHGAP39-WT. Cells were fixed and immunostained with anti-HA antibodies to visualize ARHGAP39. Scale bar, 15 μm. White boxes indicate regions selected for magnified images shown in bottom panels. Scale bar, 5 μm. White arrows indicate two representative filopodia used for line scans in (D). (D) Intensity profiles of ARHAGP39 (green) and mDia2 (red) start at the tip of the filopodia. (E and F) mDia2 depletion efficiency. (E) HeLa cells were transfected with control or mDia2 siRNA oligos for 48 h before transfection with GFP-Cdc42-Q61L (active Cdc42). Cells were labelled with phalloidin to detect F-actin. Scale bar, 10 μm. (F) Quantification of the percentage of Cdc42-expressing cells with filopodia. n ≥ 72 cells for each condition. (G and H) ARHGAP39 still localizes to filopodia in cells depleted in mDia2. (G) HeLa cells depleted in mDia2 for 48 h were transfected with GFP-Myo10 (red) and HA-ARHGAP39-WT (green). Cells were fixed and labelled with anti-HA antibodies and phalloidin to detect ARHGAP39 and F-actin, respectively. Scale bar, 15 μm. White box indicates the region selected for magnified images shown in Figure 3. (H) Quantification of the percentage of filopodia with ARHGAP39 at the tips in mDia2-depleted cells expressing Myo10. n = 166 and 605 filopodia in GFP and GFP-Myo10 expressing cells, respectively. Data are representative of two (A, B) or three (C-H) independent experiments and are presented as mean ± SEM. Statistical analysis was performed using two-tailed Student’s *t*-test. *p < 0.05; **p < 0.01 and *\*\*\**p < 1×10^−14^.

**Figure S4. Related to Figure 4.**
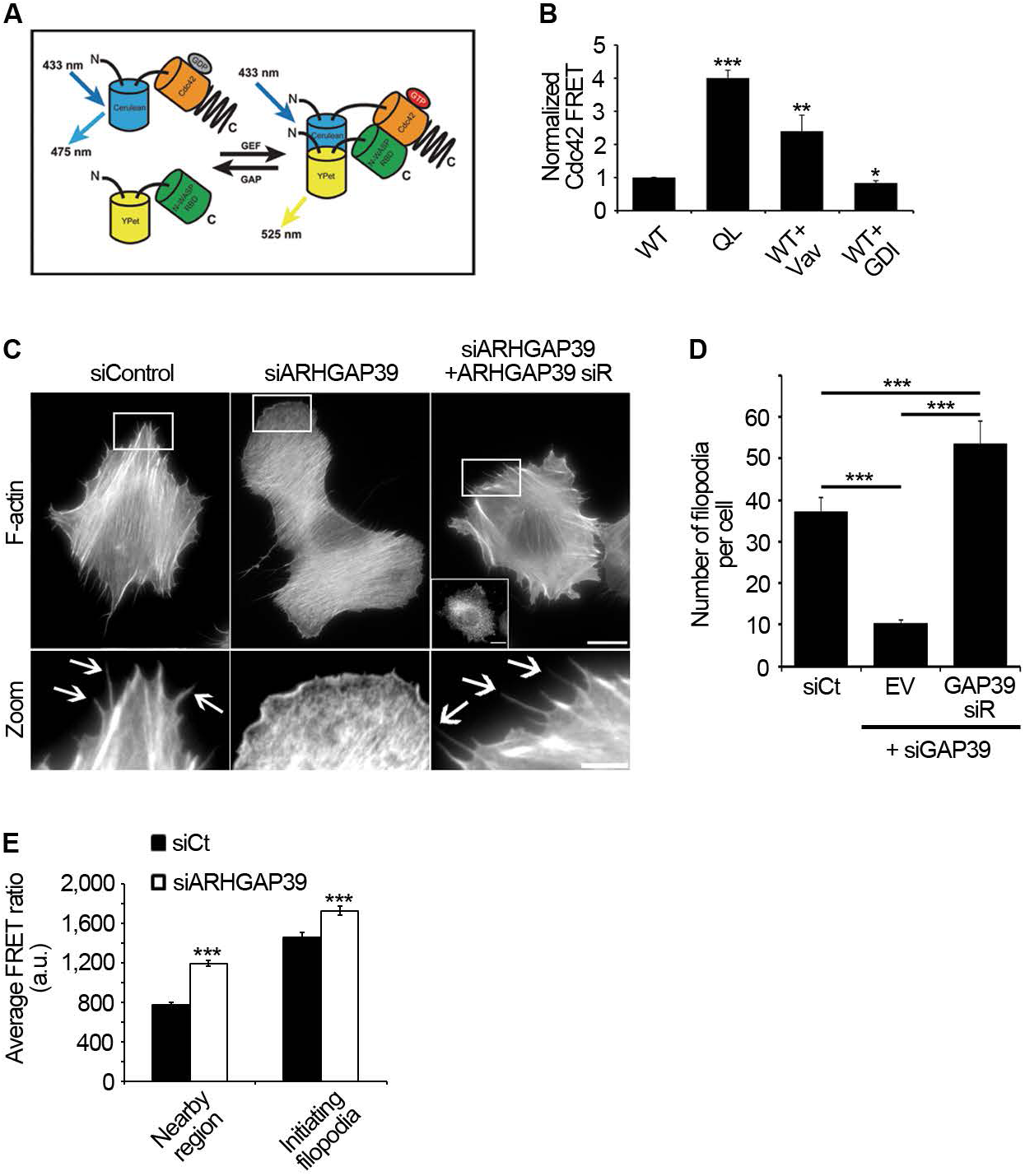
Cdc42 activity and filopodia formation are regulated by ARHGAP39. (A and B) Cdc42 biosensor design and activity. (A) Schematic representation of the intermolecular FRET Cdc42 biosensor. GTP: guanosine triphosphate, GDP: guanosine diphosphate, Cerulean: cyan fluorescent protein (CFP), YPet: yellow fluorescent protein. (B) Cdc42 activity of cell suspensions, measured in 293 cells expressing Cdc42 FRET biosensor WT or active mutant Q61L, with or without Vav2 or GDI. n = 4 (Q61L), n = 10 (Vav2) and n = 6 (GDI) independent experiments. (C and D) ARHGAP39 regulates filopodia formation. (C) HeLa cells were transfected with 20 nM of control or ARHGAP39 siRNA oligos for 48 h before transfection with HA empty vector or HA-ARHGAP39 siRNA resistant (GAP39 siR). Cells were labelled with phalloidin to detect F-actin. Scale bar, 15 μm. Inset in top panel shows expression of HA-ARHGAP39 siR detected with anti-HA antibodies. White boxes indicate regions selected for magnified images shown in bottom panel. Scale bar, 5 μm. White arrows indicate filopodia. (D) Quantification of the number of filopodia per cell. n ≥ 58 cells. (E) Cdc42 activation at the cell periphery prior to filopodium formation is increased upon ARHGAP39 depletion. At membrane regions where single filopodium formed, FRET ratio intensity in a region incorporating the base of the filopodium was measured one frame prior to filopodia formation and compared with FRET ratio intensity in a nearby region of plasma membrane of similar size. n = 46 and n = 25 filopodia in siControl and siARHGAP39, respectively. Data are representative of three independent experiments (C-E). Data are presented as mean ± SEM. Statistical analysis was performed using two-tailed STudent’s *t*-test. *\**p < 0.05; *\*\**p < 0.01 and *\*\*\**p < 0.001.

**Figure S5. Related to Figure 5.**
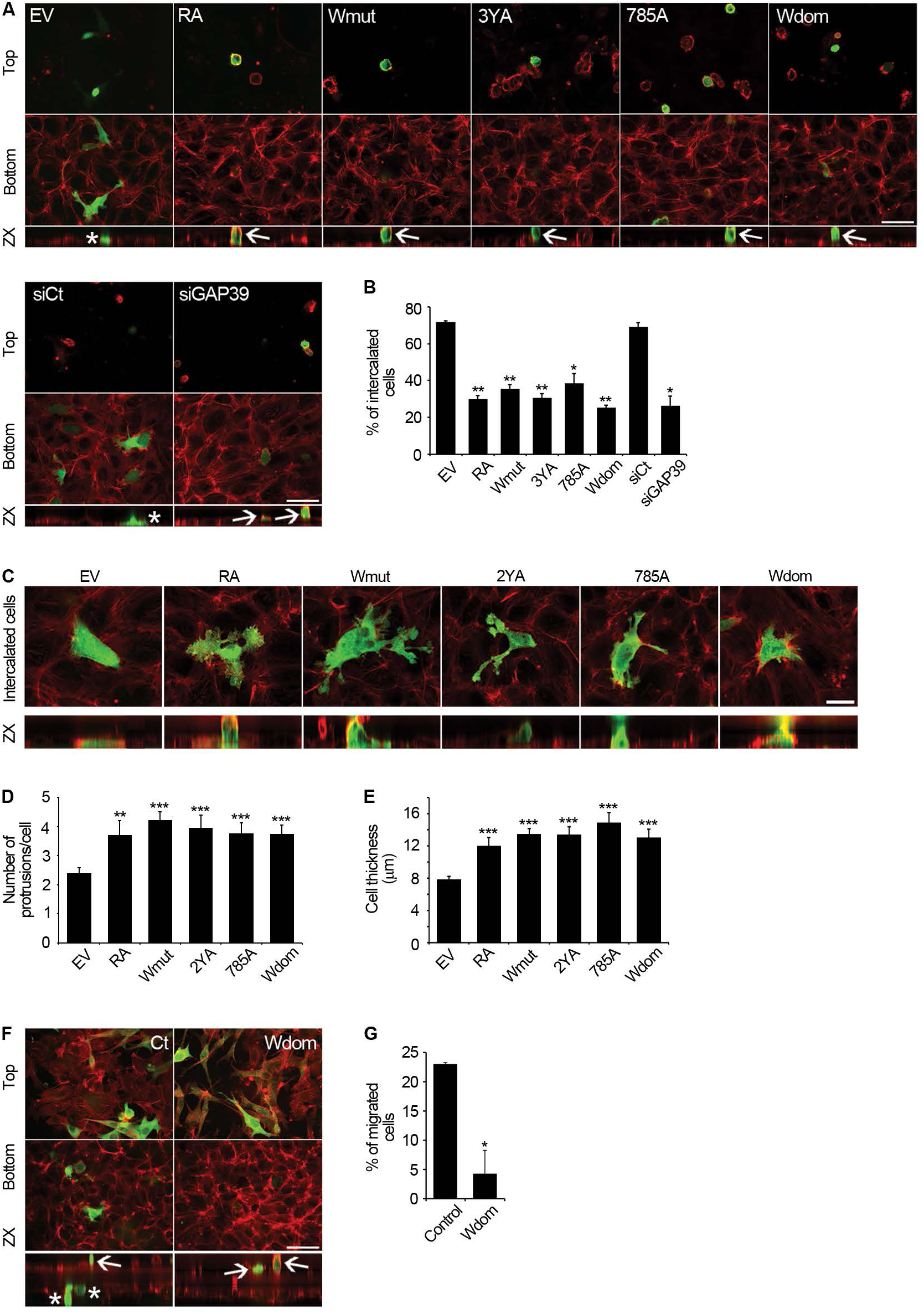
ARHGAP39 regulates cancer cell migration through endothelial cell monolayers. (A and B) Images of top and bottom of endothelial cell monolayer incubated with cancer cells transfected with GFP empty vector (EV), HA-ARHGAP39 -RA, -Wmut, −3YA, −785A, HA-Wdom, siRNA control or siRNA ARHGAP39 (A). Cells were allowed to interact for 180 min before being fixed and labelled for F-actin (red) and HA (green). Scale bar, 50 μm. Bottom panels are z projections of cells in top panels. White asterisks indicate cells that have intercalated through the endothelial cell monolayer. White arrows indicate cells that did not intercalate. (B) Quantification of the percentage of intercalated cells for each condition. n = 50-121 cells. *\**p < 0.05 and *\*\**p < 0.01 compared to EV. (C-E) Representative images (C) of cells that have intercalated through the endothelial cell monolayer from the assay in (A). Scale bar, 15 μm. Quantification of the number of protrusions (D) and cell thickness (E) of the cells that have intercalated in (A). *\*\**p < 0.001 and *\*\*\**p < 10^−4^ compared to EV. Note that even if some cells expressing ARHGAP mutants have intercalated, these cells present abnormal numbers of protrusions and have not properly integrated the endothelial monolayer compared to control cells. (F and G) Trans-endothelial migration assay with MDA-MB-435 cells stably expressing GFP empty vector control or GFP-ARHGAP39-Wdom (green). (F) Representative images of the top and bottom of transwells. Cells were labelled with phalloidin to detect F-actin (red). Scale bar, 50 μm. Bottom panels are confocal z projections of entire transwell. White asterisks indicate cells that have migrated through the endothelial cell monolayer. White arrows indicate cells that did not migrate. (G) Quantification of the percentage of migrated cells. n = 200 and 110 stable cells for control and ARHGAP39-Wdom, respectively. Data are representative of two (A siRNA, B siRNA, F, G) or three (A DNA, B DNA, C-E) independent experiments. Data are presented as mean ± SEM. Statistical analysis was performed using two-tailed Student’s *t*-test. *\*P* < 0.05 and *\*\*P* < 0.01 unless specified.

**Figure S6. Related to Figure 5.**
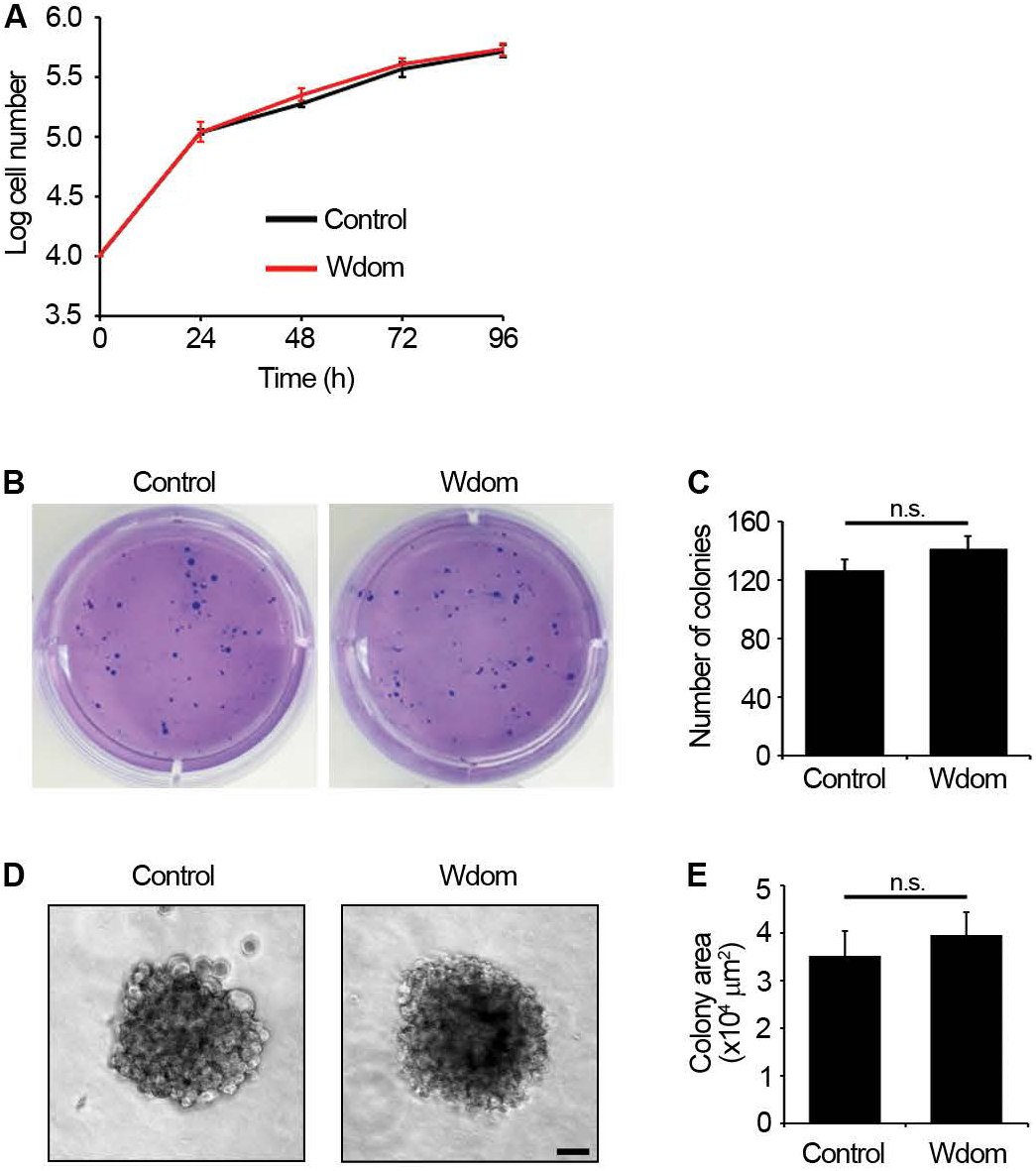
Expression of Wdom does not affect tumor cell growth and colony formation. (A) Growth curve quantification of control or ARHGAP39-Wdom-expressing stable MDA-MB-435 cells. Each measure was performed in triplicate. (B-E) Soft agar colony assay with control and ARHGAP39-Wdom-expressing stable MDA-MB-435 cells. Representative images at low resolution (B) and quantification of colony number for each cell line (C). Representative images at higher resolution (D) and quantification of colony area for each cell line (E). Scale bar, 50 μm. Each assay was performed in triplicate. Data are representative of three independent experiments. Data are presented as mean ± SEM. Statistical analysis was performed using two-tailed Student’s *t*-tests. n.s., not statistically significant compared to control.

## Video legends

**Movie S1. ARHGAP39 localizes in filopodia and its accumulation at the tip precedes filopodia retraction.** Related to Figure **3H**

HT-1080 cells expressing GFP-ARHGAP39-WT in combination with myc-Cdc42-WT were replated on collagen-coated glass coverslips before being mounted into a chamber and placed onto a temperature-controlled stage. GFP time series were acquired using a spinning disk confocal microscope. Images were acquired every 5 s. Time is shown in minutes:seconds. White arrow indicates the accumulation of ARHGAP39 at filopodia tip that precedes filopodia retraction. Scale bar, 2 μm.

**Movie S2. Cdc42 activation is detected at the leading edge and in filopodia.** Related to Figure **4A**

Representative live cell FRET ratio images of a HeLa cell transfected with the Cdc42 biosensor and plated on fibronectin-coated glass coverslips. 16 h post-transfection, cells were transferred to the microscope for imaging using a 40X objective lens. CFP, YFP and FRET images were acquired every 20 s for 15 min. Scale bar, 5 μm.

**Movie S3. Cdc42 activity correlates with filopodia dynamics.** Related to Figure **4C**

Representative live cell FRET ratio images of a HeLa cell transfected with the Cdc42 biosensor and imaged as described in Video S2. White arrows indicate Cdc42 activation that precedes initiation of filopodia. Red arrows indicate low Cdc42 activity that precedes filopodia retraction. Scale bar, 3 μm.

**Movie S4. Cdc42 activity is regulated by ARHGAP39.** Related to Figure **4F**

Representative live cell FRET ratio images of HeLa cells treated with siRNA control (left panel) or siRNA ARHGAP39 (right panel). HeLa cells were transfected with 20 nM control or ARHGAP39 siRNA oligos for 48 h before being further transfected with siRNA oligos in combination with Cdc42 biosensor. Cells were subsequently plated on fibronectin-coated glass coverslips, starved in serum-free medium for 6 h and transferred to the microscope for imaging. Cells were then stimulated with 20 ng/mL TNFα to induce filopodia formation. Cells were imaged every 20 s for 15 min. Scale bar, 5 μm.

**Movies S5 and S6. ARHGAP39 depletion affects filopodia dynamics.** Related to Figure **4K**

HeLa cells were transfected with 20 nM control (Video S5) or ARHGAP39 (Video S6) siRNA oligos for 48 h before being further transfected with siRNA oligos in combination with GFP-Myo10 to induce filopodia formation. Cells were replated on fibronectin-coated glass coverslips before being mounted into a chamber and placed onto a temperature-controlled stage. Phase-contrast time series were acquired using a 100X/1.4 NA Plan APO objective lens on a spinning disk confocal microscope. Images were acquired every 3 s for 15 min. Time is shown in s. Black arrows indicate retracting filopodia, black stars indicate collapsing filopodia. Scale bar, 2 μm.

**Movies S7 and S8. ARHGAP39 depletion decreases cell migration in 3D.** Related to Figure **5A**

HT1080 cells were transfected with 20 nM control (Video S7) or ARHGAP39 (Video S8) siRNA oligos for 24 h before being embedded into a 3D collagen gel and placed into a 4-well glass bottom chamber. The next day, chambers were placed onto a temperature-controlled stage and phase-contrast time series were acquired on our inverted microscope using a 10x/0.25 Ph1 DL Phase Contrast objective lens. Images were acquired every minute for 2.5 h. Time is shown in hours:minutes. Scale bar, 20 μm.

