## Supplemental files for "Mechanistic of Rac and Cdc42 synchronization at the cell edge by ARHGAP39-dependent signaling nodules and the impact on protrusion dynamics"

**Movies S7 and S8. ARHGAP39 depletion decreases cell migration in 3D.** Related to Figure **5A** HT1080 cells were transfected with 20 nM control (Video S7) or ARHGAP39 (Video S8) siRNA oligos for 24 h before being embedded into a 3D collagen gel and placed into a 4-well glass bottom chamber. The next day, chambers were placed onto a temperature-controlled stage and phase-contrast time series were acquired on our inverted microscope using a 10x/0.25 Ph1 DL Phase Contrast objective lens. Images were acquired every minute for 2.5 h. Time is shown in hours:minutes. Scale bar, 20 μm.

**Table 1. List of cloning primers**

| **DNA** | **Vector** | **Primers** |
|---|---|---|
| ARHGAP39 Full-length | pCMV5-HA3 | GAGAAAGCTTATGTCCCAGACG |
| | | GAGAAAGCTTCTACAGCACACCCTC |
| | pEGFP-C2 | GAGAGAATTCATGTCCCAGACGCAGGACTACG |
| | | GAGAGGATCCCTACAGCACACCCTCCATGAAG |
| | pTriEx-Flag-mCherry | GAGAAAGCTTATGTCCCAGACGCAGGACTACG |
| | | GAGAAAGCTTCTACAGCACACCCTCCATGAAG |
| ARHGAP39 Nterm (aa1-134) | pCMV5-HA3 | GAGAAAGCTTATGTCCCAGACG |
| | | GAGAAAGCTTCAGGGAGGAGCT |
| | pGEX4T1 | GAGAGAATTCATGTCCCAGACG |
| | | GAGAGAATTCCAGGGAGGAGCT |
| ARHGAP39 Nterm (aa23-97) | pBabe | GAGAGGATCCATGGGGTCGAACACTCGGTTGG |
| | | GAGAGAATTCTTAGCCCTGCGGCCGGTGCC |
| | pBabe GFP | GAGAGGATCCATGGGGTCGAACACTCGGTTGG |
| | | GAGAGGATCCTTAGCCCTGCGGCCGGTGCC |
| ARHGAP39 Cterm (aa135-1084) | pCMV5-HA3 | GAGAAAGCTTAGCACCAGCTCC |

| | | |
|---|---|---|
| | | GAGAAAGCTTCTACAGCACACCCTC |
| ARHGAP39-Y313A | | GAACCCCCCGCTCGCCGAGGAGCCCCCA |
| | | TGGGGGCTCCTCGGCGAGCGGGGGTTC |
| ARHGAP39-Y325A | | CAGGCCCCCATCGCCGATGAGCCCCCC |
| | | GGGGGGCTCATCGGCGATGGGGGCCTG |
| ARHGAP39-Y785A | | CGGGACGAGCTCGCCATCCAGCTGTGC |
| | | GCACAGCTGGATGGCGAGCTCGTCCCG |
| ARHGAP39-Wmut (W53A) | | GTGAGTGCGTGGCGGACCCGCCG |
| | | CGGCGGGTCCGCCACGCACTCAC |
| ARHGAP39-Wmut (W53A, P56A) | | GCGGACCCGGCGGCCGGCG |
| | | CGCCGGCCGCCGGGTCCGC |
| ARHGAP39-Wmut (W92A) | | GCGCACGGTGGCGCACCGGCCG |
| | | CGGCCGGTGCGCCACCGTGCGC |
| ARHGAP3-Wmut (W92A, P95A) | | GGCGCACCGGGCGCAGGGCTG |
| | | CAGCCCTGCGCCCGGTGCGCC |
| ARHGAP39-RA (R932A) | | GACAGAGGGCATCTTCGCGGTCCCTGGGGACATTG |
| | | CAATGTCCCCAGGGACCGCGAAGATGCCCTCTGTC |
| Myo10-Myth-FERM (aa1486-2058) | pGEX4T1 | GAGA GCGGCCGCAT ACCCGGTGGTCCAGTGCC |
| | | GAGA GCGGCCGC TCACCTGGAGCTGCCCTG |
| Wave2 Full-length | pCDNA3-HA | GAGAGGTACCATGCCGTTAGTA |
| | | GAGAGGTACCTTAATCGGACCAGTC |
| Wave2 Nterm (aa1-299) | pCDNA3-HA | GAGAGGTACCATGCCGTTAGTA |
| | | GAGAGGTACCTGCATCCACGTT |
| Wave2 Cterm (aa420-498) | pCDNA3-HA | GAGAGGTACCACCACCAAGCCC |
| | | GAGAGGTACCTTAATCGGACCAGTC |
| Wave2 poly-proline region (aa300-419) | pCDNA3-HA | GAGAGGTACCAGTAGCTATCCG |
| | | GAGAGGTACCATCAGAAAGCGG |
| mDia2 Full-length | pCDNA3-HA | GAGAGGTACCATGGAACGGCAC |
| | | GAGAGGTACCTTATAAAGCTCGTAA |
| mDia-deltaDAD (aa1-1057) | pCDNA3-HA | GAGAGGTACCATGGAACGGCAC |
| | | GAGAGGTACCATCACCCTCAGT |
| mDia2 Nterm (aa1-514) | pCDNA3-HA | GAGAGGTACCATGGAACGGCAC |

| | | |
|---|---|---|
| | | GAGAGGTACCCTTGTAAAGTTC |
| mDia2 Cterm (aa689-1193) | pCDNA3-HA | GAGAGGTACCCAACAAAAAGAG |
| | | GAGAGGTACCTTATAAAGCTCGTAA |
| mDia2 FH1 domain (aa506-696) | pCDNA3-HA | GAGAGGTACCGAAGAGAAAGCA |
| | | GAGAGGTACCCTCTTCTCTTCT |

**Figure S1. Related to Figure 1.**

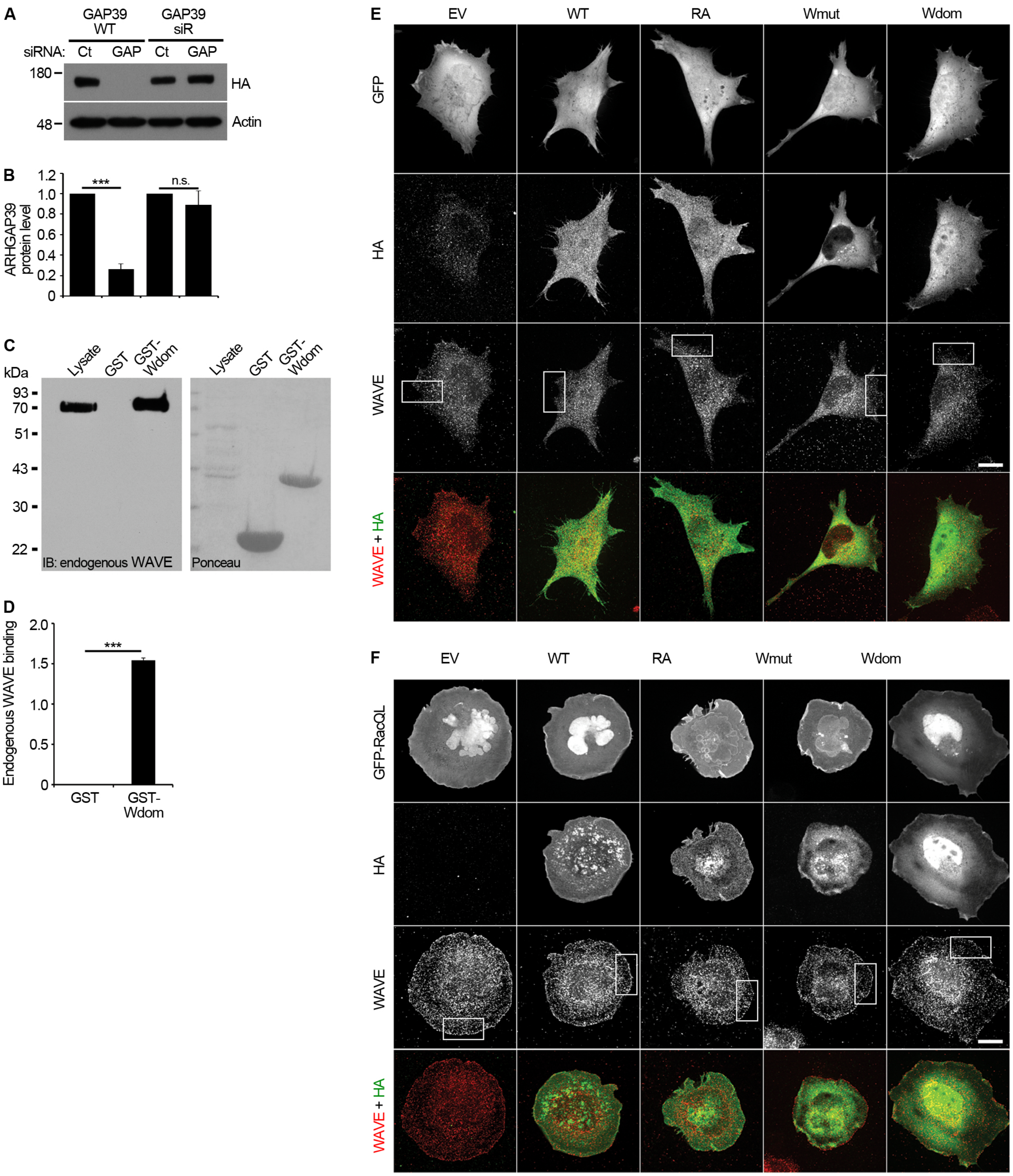

**Figure S2. Related to Figure 2.**

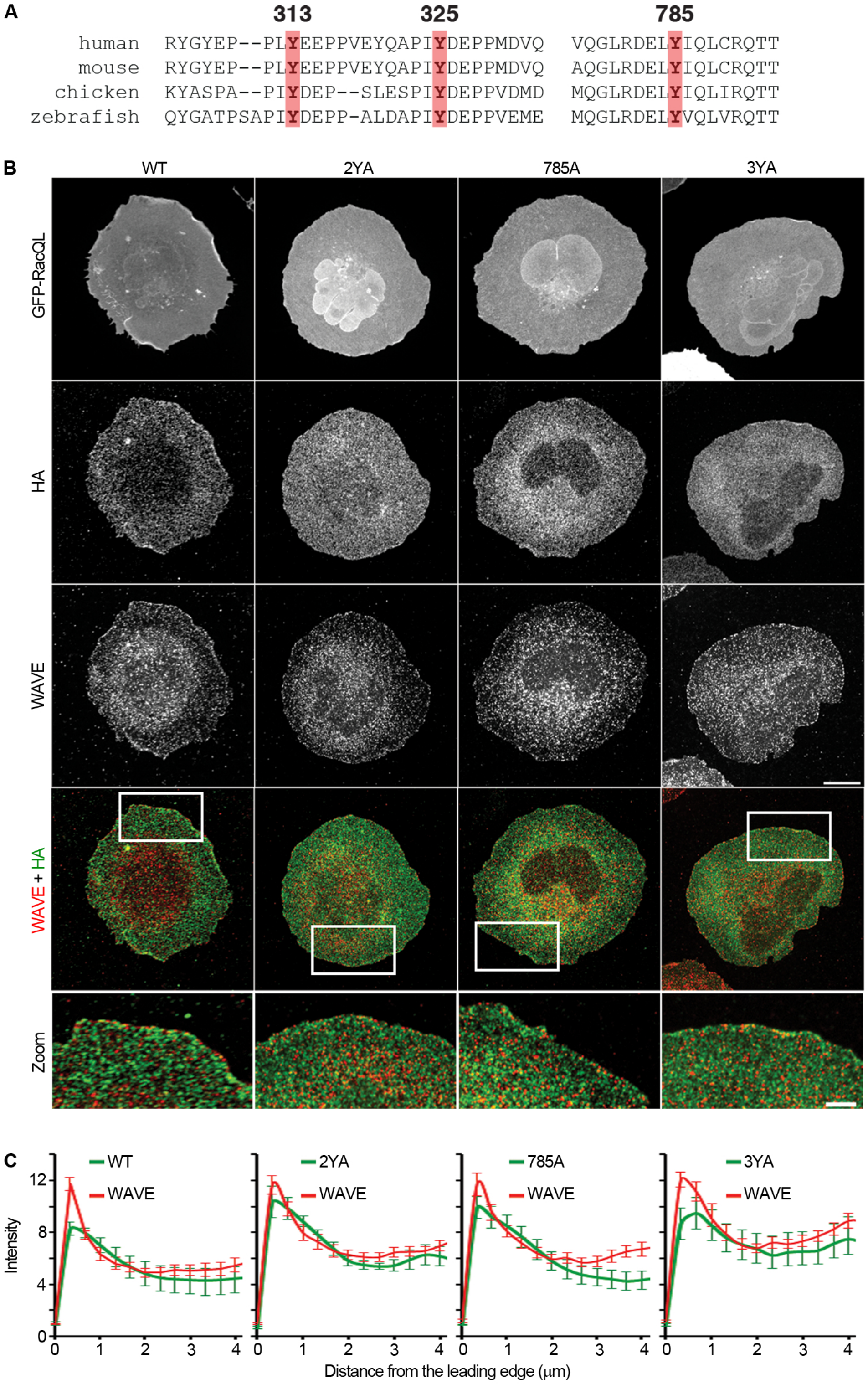

**Figure S3. Related to Figure 3.**

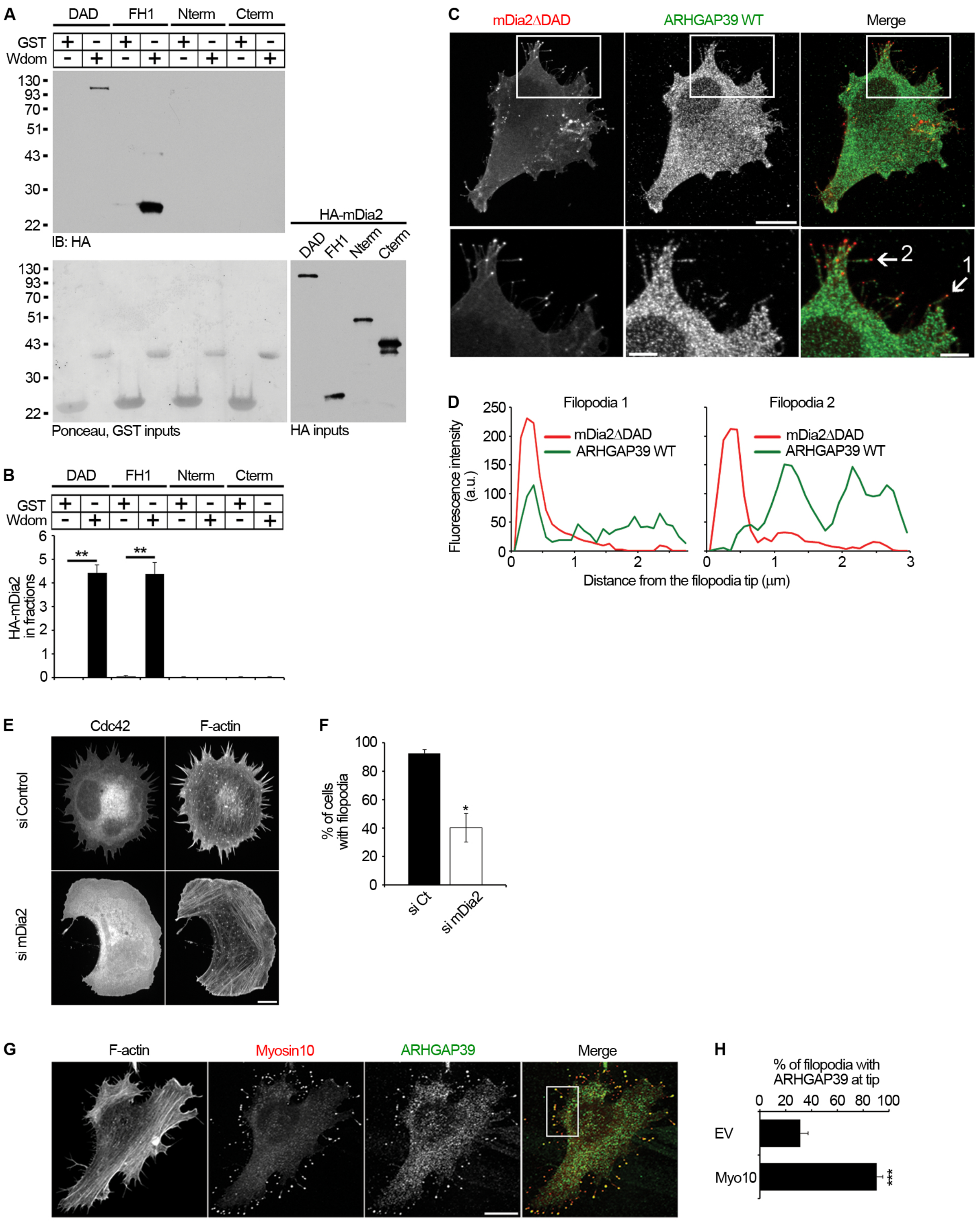

**Figure S4. Related to Figure 4.**

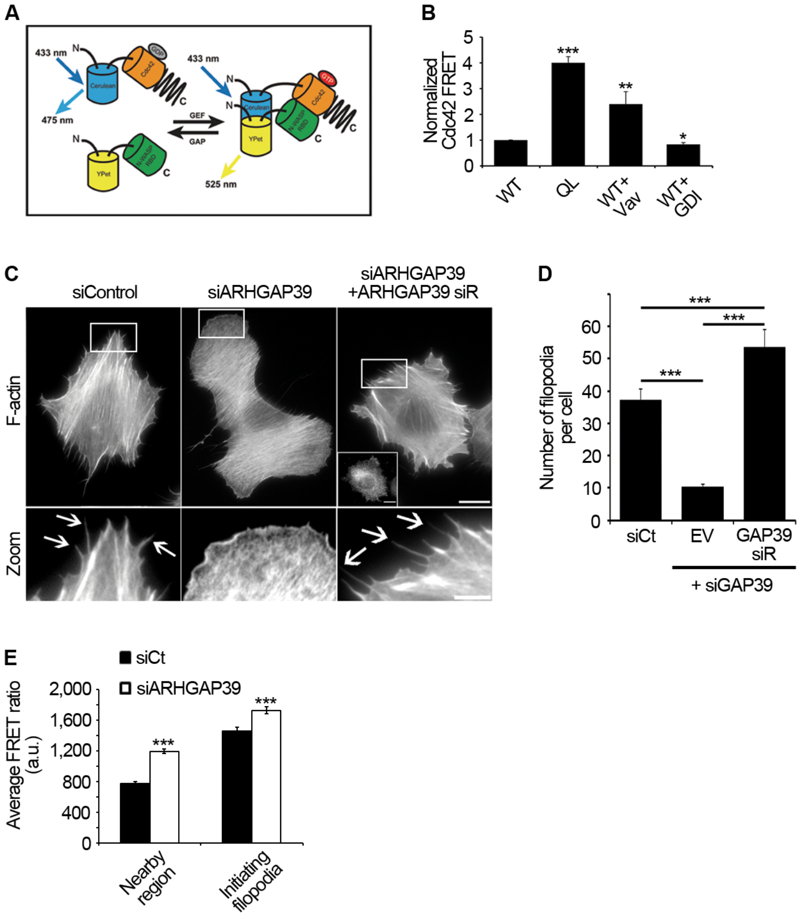

**Figure S5. Related to Figure 5.**

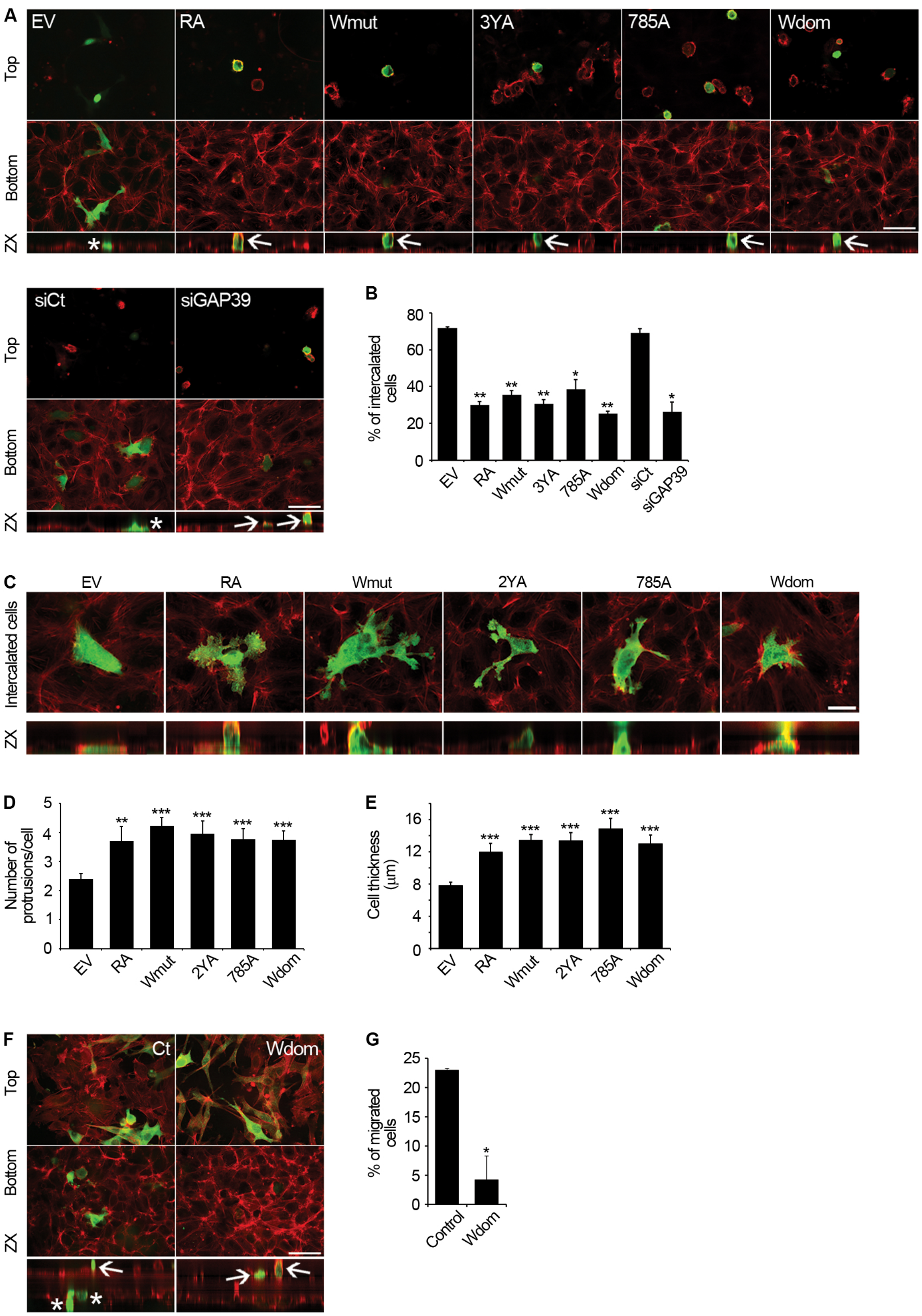

**Figure S6. Related to Figure 5.**

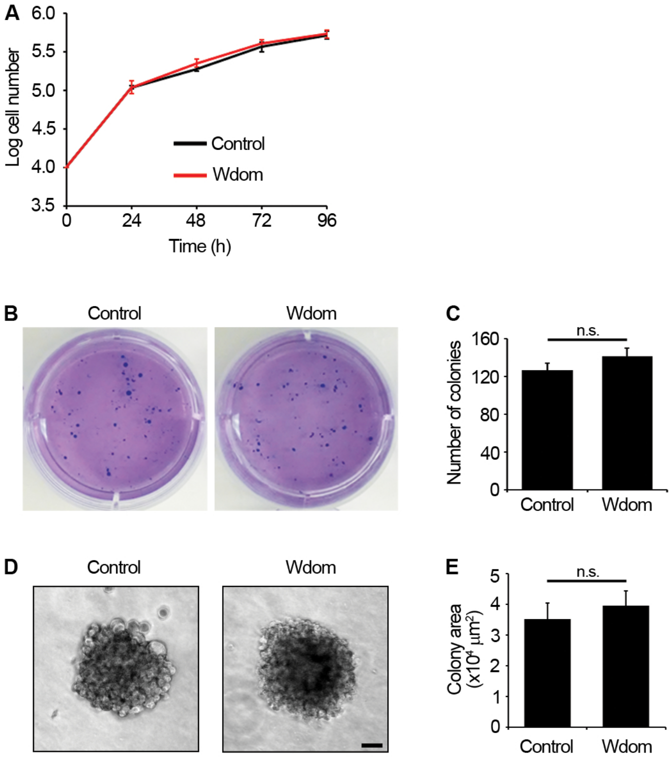
